# RPG acts as a central determinant for infectosome formation and cellular polarization during intracellular rhizobial infections

**DOI:** 10.1101/2022.06.03.494689

**Authors:** Beatrice Lace, Chao Su, Daniel Invernot Perez, Marta Rodriguez-Franco, Tatiana Vernié, Morgane Batzenschlager, Sabrina Egli, Cheng-Wu Liu, Thomas Ott

## Abstract

Host-controlled intracellular accommodation of nitrogen-fixing bacteria is essential for the establishment of a functional Root Nodule Symbiosis (RNS). In many host plants, this occurs via transcellular tubular-structures (infection threads - ITs) that extend across cell layers via polar tip- growth. Comparative phylogenomic studies have identified RPG (RHIZOBIUM-DIRECTED POLAR GROWTH) among the critical genetic determinants for bacterial infection. In *Medicago truncatula*, RPG is required for effective IT progression within root hairs but the cellular and molecular function of the encoded protein remain elusive. Here, we show that RPG resides in the protein complex formed by the core endosymbiotic components VAPYRIN (VPY) and LUMPY INFECTION (LIN) required for IT polar growth, co- localizes with both VPY and LIN in IT tip- and perinuclear-associated puncta of *M. truncatula* root hairs undergoing infection and is necessary for VPY recruitment to puncta. Fluorescence Lifetime Imaging Microscopy (FLIM) of phosphoinositide species during bacterial infection revealed that functional RPG is required to sustain strong membrane polarization at the advancing tip of the IT. In addition, loss of RPG functionality alters the cytoskeleton-mediated connectivity between the IT tip and the nucleus and affects polar secretion of the cell wall modifying enzyme NODULE PECTATE LYASE (NPL). Our results integrate RPG into a core host machinery required to support symbiont accommodation, suggesting that its occurrence in plant host genomes is essential to co-opt a multimeric protein module committed to endosymbiosis to sustain IT-mediated bacterial infection.

## INTRODUCTION

Legumes have evolved the capacity to maintain a mutualistic association with endosymbiotic nitrogen-fixing rhizobia, hosting them in specialized lateral root organs, the nodules, where bacteria convert atmospheric nitrogen (N2) to ammonia (NH4^+^) and release it to the plant in exchange for carbohydrates. By granting them access to an additional source of nitrogen, the Root Nodule Symbiosis (RNS) allows plants to overcome nitrogen limitation occurring in terrestrial ecosystems and agriculturally used sites. Thus, the transfer of this ability to cereals has been identified as one possible strategy to reduce the use of environmentally harmful and expensive industrial fertilizers (S.E. Bloch et al., 2020; Mus et al., 2016; Oldroyd & Dixon, 2014; Pankievicz et al., 2019; Rogers & Oldroyd, 2014). Such an engineering approach however, requires an in depth understanding of the molecular machineries allowing rhizobial infections of legume roots.

In most cases, this host-controlled process requires a molecular dialogue between the partners, which involves the perception of bacterial lipo-chitooligosaccharide (LCOs) molecules called Nod factors (NFs) by plasma membrane-resident receptor complexes including the *Medicago truncatula* LysM receptor-like kinase NFP and its *Lotus japonicus* ortholog NFR5 (Limpens et al., 2003; Radutoiu et al., 2003). Symbiont recognition triggers the onset of a signaling cascade involving the generation and decoding of nuclear-associated calcium oscillations, which mediates symbiosis-related gene expression (Singh et al., 2014). Interestingly, the same pathway is activated by perception of diffusible signals produced by arbuscular mycorrhizal (AM) fungi, including both LCOs and long-chain chitooligosaccharides (COs), although these are perceived by a different combination of receptors (Feng et al., 2019; Sun et al., 2015). This common symbiosis signaling pathway shares a number of genetic components including the machinery to decode the specific calcium signature (Kistner et al., 2005) and few downstream factors such as VAPYRIN (VPY), a Major Sperm Domain- and ankyrin-repeat-containing protein, that is required for both symbiotic interactions (Murray et al., 2011; Pumplin et al., 2010). Although some structural features are shared between the AM symbiosis and RNS during intracellular colonization, others are remarkably different. A whole set of those RNS-specific responses is controlled by the master regulator NIN, a transcription factor orchestrating the transcriptional reprogramming of cells required to sustain bacterial accommodation and concomitant cell proliferation to induce the formation of a nodule primordium (Schauser et al., 1999; C. W. Liu et al., 2019a; Vernié et al., 2016). This spatial and temporal coordination of infection and nodule organogenesis represents a hallmark of RNS (Oldroyd & Downie, 2008).

Intracellular invasion of rhizobia is uniquely enabled by a membrane-confined transcellular tunnel called the infection thread (IT). In many legumes such as *M. truncatula* and *L. japonicus* IT formation is initiated from an infection chamber (IC) (Fournier et al., 2015) that forms upon curling of a growing root hair and subsequent entrapping of rhizobia (Callaham & Torrey, 1981). Repolarization of targeted secretion towards this compartment enables local cell wall modifications (e.g. by secretion of the NODULE PECTATE LYASE (NPL) to the IC (Liu et al., 2019a)) leading to its radial expansion (Fournier et al., 2015) and a highly confined local invagination of the plasma membrane, which marks the onset of IT initiation (Gage, 2004). The IT then extends through the trichoblast and subsequently progresses transcellularly through the root cortex (Libbenga & Harkes, 1973). IT maintenance and progression requires a highly coordinated interplay between the migrating nucleus, a cytosolic column ahead of the IT, actin and dense microtubule arrays (Fournier et al., 2015; Perrine- Walker et al., 2014; Qiu et al., 2015; Timmers et al., 1999; Yokota et al., 2009). IT propagation in underlying cortical tissues is supported by repolarization of these cells, forming pre-infection thread (PIT) structures (Van Brussel et al., 1992). This ensures guidance of the symbionts towards the newly divided cortical cells constituting the core of the nodule primordium. There, rhizobia are released from the ITs into the host cell, where they differentiate into nitrogen-fixing bacteroids (Vasse et al., 1990).

Over the last decades, reverse and forward genetics have identified several genes required for bacterial infection (Roy et al., 2020; Tsyganova et al., 2021). Among these is *RHIZOBIUM- DIRECTED POLAR GROWTH (RPG)*, which is specifically expressed upon rhizobial inoculation and encodes a protein of unknown function (Arrighi et al., 2008). A loss of RPG in *M. truncatula* results in the development of aberrant ITs and poorly colonized nodules (Arrighi et al., 2008). Nuclear localization of RPG when heterologously over-expressed in *Nicotiana benthamiana* leaf epidermal cells suggested a putative role as a transcription factor (Arrighi et al., 2008), but its cellular and molecular function in *M. truncatula* have not been understood in detail. The protein, however, recently re-gained great attention, when two independent comparative phylogenomic studies identified *NFP*, *NIN* and *RPG* as the only genes that have been consistently lost in non-nodulating species belonging to the Fagales/Fabales/Cucurbitales/Rosales (FaFaCuRo) clade (Griesmann et al., 2018; van Velzen et al., 2018). This evolutionary pattern linking these genes and the ability to form RNS makes them prime candidates for engineering symbiotic nitrogen fixation in cereals.

Here, we demonstrate that RPG is an essential component of the infectosome complex (Liu et al., 2019a; Roy et al., 2020). This multi-protein assembly associates with the very tip of growing ITs and with the nuclear periphery. A loss-of function mutation in *RPG* entirely blocks the recruitment of VPY to the infectosome. Furthermore, a highly confined membrane polarity domain at the IT tip, cytoskeleton connectivity between this site and the nucleus and targeted secretion to the IT are aberrant in an *rpg* mutant. Thus, we propose that RPG provides specificity to a conserved cellular machinery that enables rhizobial accommodation.

## RESULTS

### *RPG* controls development of infection thread structures in *M. truncatula*

An ethyl methanesulfonate (EMS)-induced mutant of *M. truncatula* accession A17 (hereafter named *rpg-1,* Figure 1A) was previously reported to develop enlarged ITs and poorly colonized nodules (Arrighi et al., 2008), indicating that *RPG* is required for rhizobial infection. In order to genetically confirm this function, we isolated a second independent, homozygous mutant allele from the *M. truncatula Tnt1* transposon insertion collection in the R108 accession carrying an insertion at 948bp downstream of the start codon (NF11990, hereafter named *rpg-2*, Figure 1A). We phenotypically analyzed both mutant alleles in parallel. Confocal imaging of infected root hairs 35 days post- inoculation (dpi) with a fluorescent GFP-tagged strain of the compatible symbiont *S. meliloti* (*S. meliloti*-GFP) revealed similar alterations of IT morphology in both *rpg* mutants, with ITs showing a distinctive bulbous and thick appearance compared to straight and thin ITs of their corresponding wild-types (Figure 1B). To better characterize and quantitatively evaluate this peculiar IT phenotype, we scored the diameter of the tube at three different points along the IT (proximal to the infection chamber, intermediate and distal, Figure 1C). While the diameter size of ITs developed by the wild- type had little variation (Figure 1D), reflecting the tight regulation exerted by the host over development of this tubular structure, the diameter of ITs formed in roots of the *rpg* mutants was highly variable and significantly wider at all measure points (Figure 1D). Altogether, this indicates loss of stringent host-controlled IT maintenance in these mutants.

**Figure 1.**
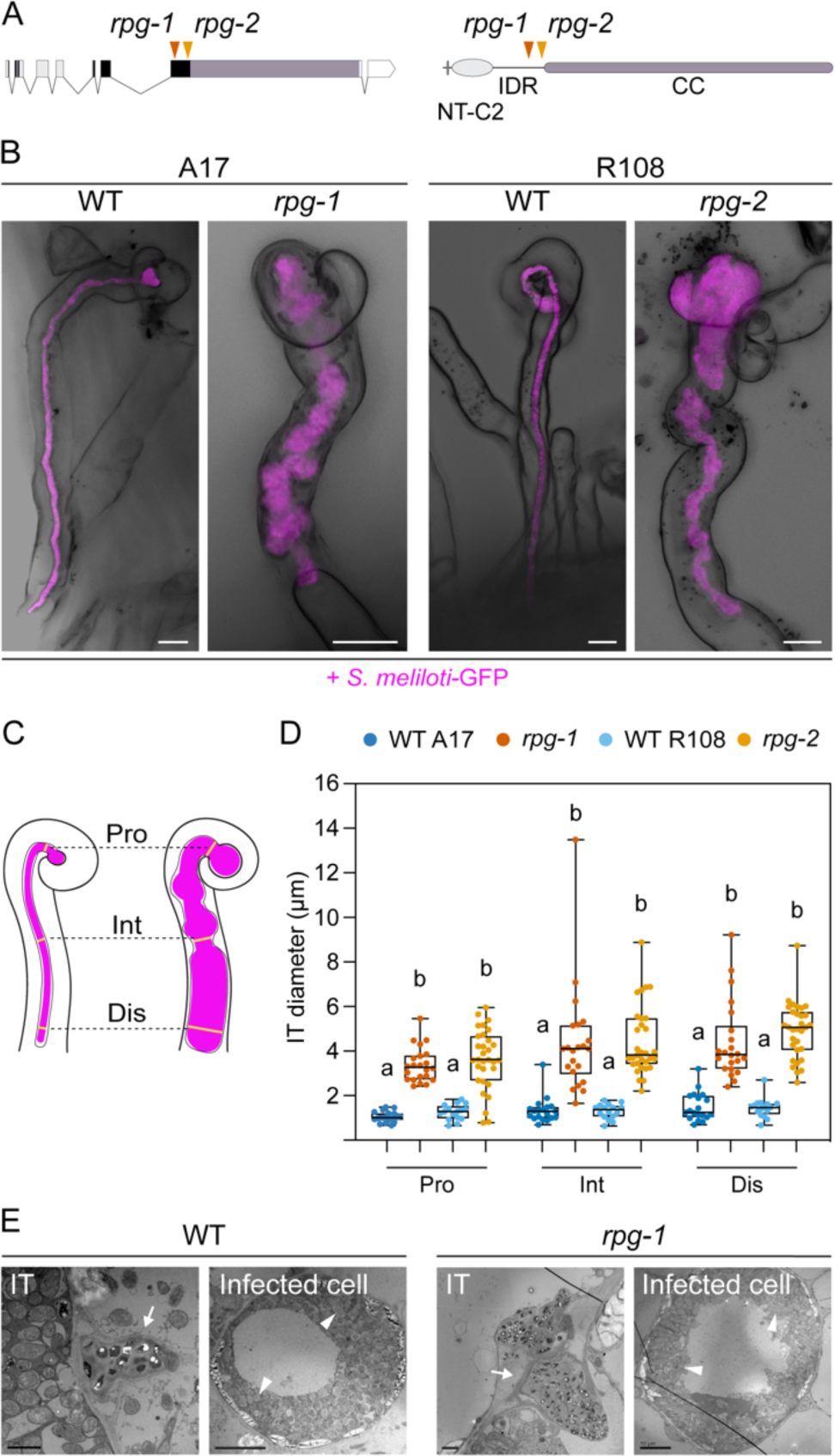
*RPG* is required for the maintenance of infection thread morphology. **(A**) Gene (left) and protein (right) structure of RPG showing the position of the mutations of two different mutant alleles (*rpg-1*, Arrighi *et al*., 2008, orange arrowhead; *rpg-2*, this study, yellow arrowhead). Both mutations map in the third exon of the *RPG* gene corresponding to the disordered region of the protein located upstream of the coiled-coil domain. NT-C2 = N-terminal C2 domain; IDR = intrinsically disordered region; CC = coiled-coil domain. (**B)** Representative confocal images of aberrant ITs formed 35 dpi within root hairs of *rpg-1* and *rpg-2* roots compared to thin and elongated ITs of the corresponding wild-types (WT). Images are overlaid of intensity projections of fluorescence and brightfield channels. *S. meliloti*-GFP is shown in magenta. Scale bars = 10 µm. (**C)** Schematic visualization of the method used to quantify morphological IT defects. The IT diameter was measured on the fluorescent channel at three different points along the IT length: in proximity of the infection chamber (Pro), in the intermediate part of the IT (Int) and in the distal part of the IT (Dis). (**D**) IT diameters scored on roots of *rpg* mutants and corresponding wild-types at the three measured points shown in **C**. In the box plot, the top and bottom of each box represents the 75th and 25th percentiles, the middle horizontal bar indicates the median and whiskers represent the range of minimum and maximum values. Letters indicate statistically significant differences according to Kruskal-Wallis multiple comparison analysis followed by Dunn’s post-hoc test. The number of plants analyzed per genotype is 4 (WT A17), 5 (*rpg-1*); 4 (WT R108); 4 (*rpg-2*) with at least 2 ITs per plant being analyzed. **(E)** TEM sections obtained from WT and *rpg-1* nodules showing IT structures (arrows) with thicker appearance in *rpg-1* compared to WT. The organization of infected cells and symbiosome (arrowheads) morphology are similar in the two genotypes. Scale bars (from left to right) = 2 µm, 10 µm, 2 µm, 10 µm.

Despite their defective growth, ITs of both *rpg* mutants occasionally penetrated cortical cell layers. To examine the morphology of cortical ITs with cellular resolution, we cleared and imaged roots at 14 dpi with *S. meliloti*-mCherry and stained them with the cell wall stain Calcofluor white to highlight the cell borders. Similar to what we observed in root hairs, ITs within cortical cells appeared consistently enlarged and bulbous in roots of both *rpg* mutants alleles compared to their wild-type genotypes, indicating that *RPG* is required to control IT development in both epidermal and cortical cell layers (Figure 1-figure supplement 1A).

We next compared the nodulation capacity of the mutant alleles by quantifying the number and type of nodules formed 21 dpi with *S. meliloti-*LacZ. We found that nodular structures formed on roots of both *rpg* mutant alleles, but the total number of infected nodules was significantly reduced compared to their corresponding wild-types. In addition, *rpg* mutants failed to develop fully elongated nodules at this time point (Figure 1-figure supplement 1B). This further supports the hypothesis that *RPG* is not required to initiate nodule organogenesis but is essential for efficient infection and full colonization. Nodulation defects were more pronounced in *rpg-2* than in *rpg-1*, which well correlates with previous findings showing that a mutation within the infection-related gene *LIN,* which is required for IT growth, affects more severely nodule development when occurring in the genetic background R108 compared to A17 (Guan et al., 2013).

To assess the relevance of *RPG* for bacterial release inside nodules, we examined ultra-thin sections of infected nodules formed on *rpg-1* and A17 WT via transmission electron microscopy (TEM). TEM micrographs of nodule infected cells revealed larger IT structures present in nodule tissues of *rpg-1* compared to WT but WT-like cell organization and symbiosome morphology (Figure 1E), indicating that normal IT development in nodules requires *RPG*, but bacterial release and differentiation are not directly dependent on this gene.

Altogether these data genetically confirm that *RPG* is essential to sustain efficient bacterial infection in *M. truncatula* and clearly support a role in controlling proper assembly and progression of infection thread structures in host root tissues.

### The coiled-coil domain of RPG is necessary for infection thread maintenance

To further understand how RPG mediates IT maintenance, we analyzed the molecular determinants required to mediate the functionality of the encoded protein during infection of root hairs.

*RPG* encodes a 1255 aa long protein with an N-terminal segment containing a putative nuclear localization signal (NLS) and an N-terminal C2 domain (NT-C2) predicted to mediate membrane tethering (Zhang et al., 2010), linked by a serine-rich intrinsically disordered region (IDR) to a long alpha-helical C-terminal extension with several predicted coiled-coil (CC) regions possibly mediating oligomerization (Figure 2A). A similar structural organization and high degrees of sequence conservation of the NT-C2 domain and the N-terminal part of the coiled-coil region were found in several homologous proteins (Arrighi et al., 2008, Figure 2-figure supplement 1). However, since any of these proteins remained uncharacterized at the functional and molecular level, the contribution of the structural features to protein functionality are unknown.

**Figure 2.**
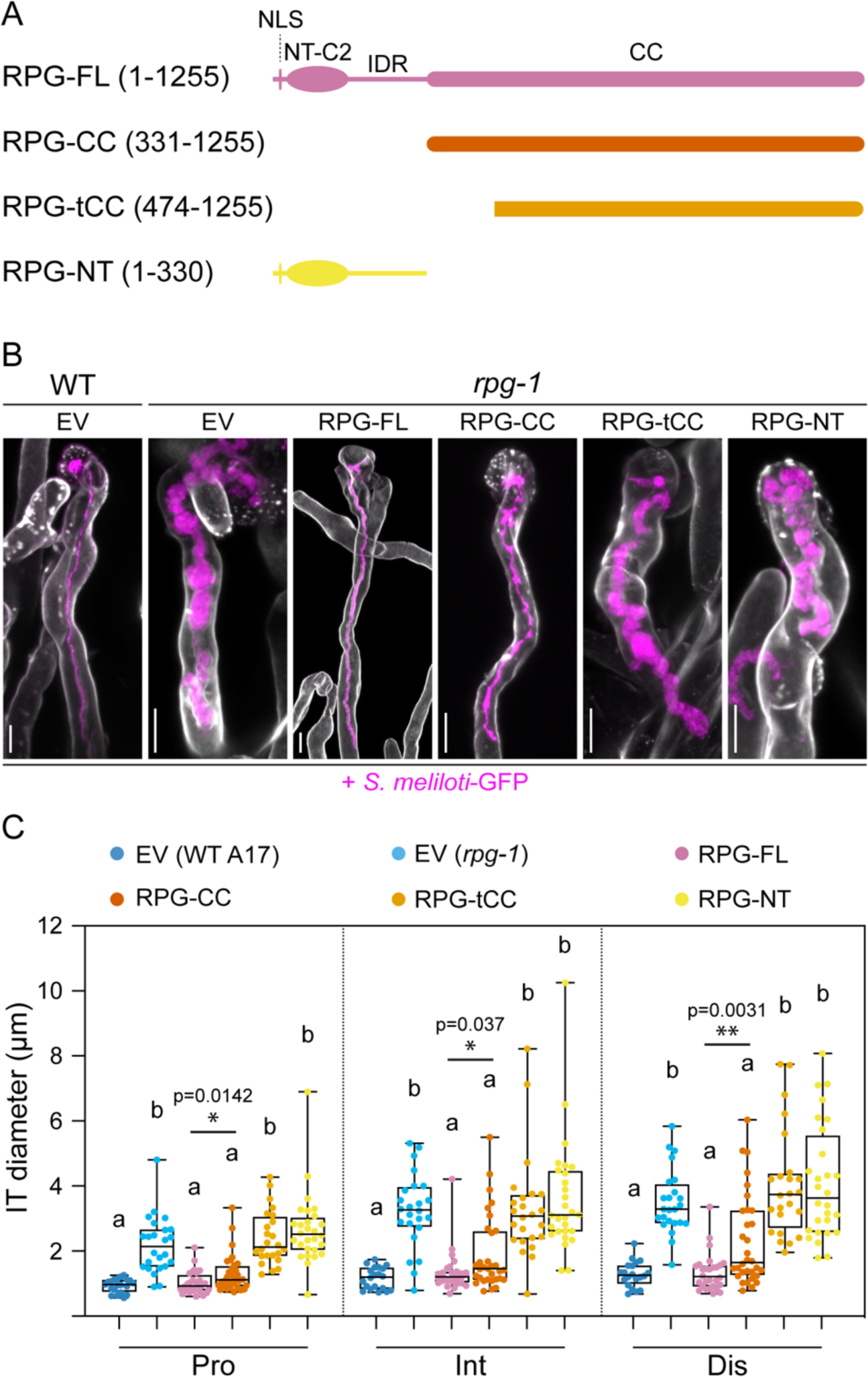
The coiled-coil domain of RPG is necessary to restore WT-like infection thread morphology in *rpg-1* root hairs. **(A)** Schematic representation of the RPG full-length protein and deletion derivatives used for complementation assays. Numbers indicate amino acids included in each fragment. NLS = nuclear localization signal; NT-C2 = N-terminal C2 domain; IDR = intrinsically disordered region; CC = coiled-coil domain. **(B)** Representative confocal images of ITs developed within root hairs of WT and *rpg-1* transgenic roots expressing the different constructs 21 dpi with *S. meliloti-* GFP (magenta). Cell walls were stained with Calcofluor white (white). Images are merge of intensity projections of fluorescent channels. EV = empty vector. Scale bars = 10 µm **(C)** IT diameters scored at the three different points schematically represented in Figure 1C. Data are presented as box plots where the top and bottom of each box represents the 75th and 25th percentiles, the middle horizontal bar indicate the median and whiskers represent the range of minimum and maximum values. Letters indicate statistically significant differences according to Kruskal-Wallis multiple comparison analysis followed by Dunn’s post-hoc test. A Mann-Whitney test was performed to compare RPG-FL and RPG-CC with *p*-values <0.05 (*), <0.01 (**), < 0.001 (***). The number of composite plants analyzed per each condition is 5 (EV in WT), 6 (EV in *rpg-1*), 7 (RPG-FL), 8 (RPG-CC), 6 (RPG-tCC), 7 (RPG-NT). For each plant, 4 ITs have been analyzed.

To gain more insights into the structural requirements for RPG functionality, we generated a series of truncation derivatives corresponding to i) the alpha-helical C-terminal coiled-coil region (aa 331- 1255, RPG-CC; Figure 2A), ii) an N-terminal truncated version of the coiled-coil region, lacking the region conserved among RPG homologues (aa474-1255; RPG-tCC; Figure 2A) and iii) the N- terminal part containing the NLS, the NT-C2 domain and the disordered region (1-330; RPG-NT; Figure 2A). To further extend localization pattern analysis, we additionally produced N-terminal mCherry fusions of the full-length protein (RPG-FL; Figure 2A) and all truncated variants. Functionality of the constructs was first assessed by testing their capability to genetically complement IT morphology when expressed in *rpg-1* roots under the control of a functional native *RPG* promoter. Mutant plants expressing mCherry-RPG-FL phenocopied the WT-like IT morphology, indicating that N-terminal tagging does not impair protein functionality during IT development within root hairs (Figure 2B-C). Interestingly, we found that the sole coiled-coil region of RPG (mCherry-RPG-CC) was equally able to significantly restore the morphological defects of *rpg-1* ITs, even though with lower efficiency when directly compared to RPG-FL (Figure 2B-C). This ability was fully dependent on the N-terminal 143 aa of this region, as expression of mCherry-RPG-tCC did not rescue the *rpg-1* IT phenotype (Figure 2B-C). This suggests that major molecular determinants conferring functionality to RPG reside in its coiled-coil domain. This was further supported by the fact that the mCherry-NT variant alone did not restore normal IT growth in the *rpg-1* mutant (Figure 2B-C). On the other side, the lower complementation efficacy of mCherry-RPG-CC compared to mCherry-RPG- FL (Figure 2C) clearly indicates the relevance of the N-terminal region for full RPG functionality and further suggests that the NLS, the NT-C2 domain and the disordered region might constitute an important regulatory unit required to fine-tune the protein function or regulate its stability.

### RPG resides in the VPY-LIN-EXO70H4 protein complex and modulates VPY recruitment to puncta

To analyze RPG localization patterns, we performed live-cell imaging of WT *M. truncatula* roots (A17) expressing mCherry-RPG-FL under the control of the native *RPG* promoter. Following rhizobial inoculation, the RPG protein consistently coalesced into few discrete cytoplasmic puncta that were visible in both uninfected and infected root hairs imaged 4-5 dpi (Figure 3A). In uninfected root hairs, such puncta were found in the vicinity of the tip (uRH; Figure 3A) while in curled root hairs at the early stage of infection, these puncta located in close proximity to the infection chamber (IC), often adjacent to the entrapped bacteria (IC; Figure 3A). In root hairs harboring growing ITs, RPG-labelled puncta were systematically present in the perinuclear cytoplasm and associated with the tip of the extending IT (IT; Figure 3A). Closer inspection showed that IT tip-associated puncta encased the growing IT tip ahead of progressing bacteria (Figure 3-figure supplement 1A). When ITs exhibited multiple branches, RPG-labelled puncta associated to the tip of each branch (Figure 3- figure supplement 1B). Occasionally, RPG signal was observed in the nucleus and the cytoplasm of infected root hairs, but its intensity was markedly weaker compared to the signal intensity detected in the puncta (Figure 3-figure supplement 1C-D). These data indicate that RPG preferentially accumulates in cytoplasmic puncta associated to infection sites during bacterial accommodation within root hairs. This is in contrast to the exclusive nuclear localization previously reported for RPG when being ectopically expressed in *N. benthamiana* leaves (Arrighi et al., 2008). While we also observed nuclear localization of RPG upon constitutive, heterologous over-expression in *N*. *benthamiana* leaves, we additionally detected RPG signal accumulating in the nucleolus and in perinuclear-localizing structures (Figure 3-figure supplement 2). Such accumulation of RPG in perinuclear structures was markedly enhanced by addition of the silencing suppressor p19 (Figure 3-figure supplement 2).

**Figure 3.**
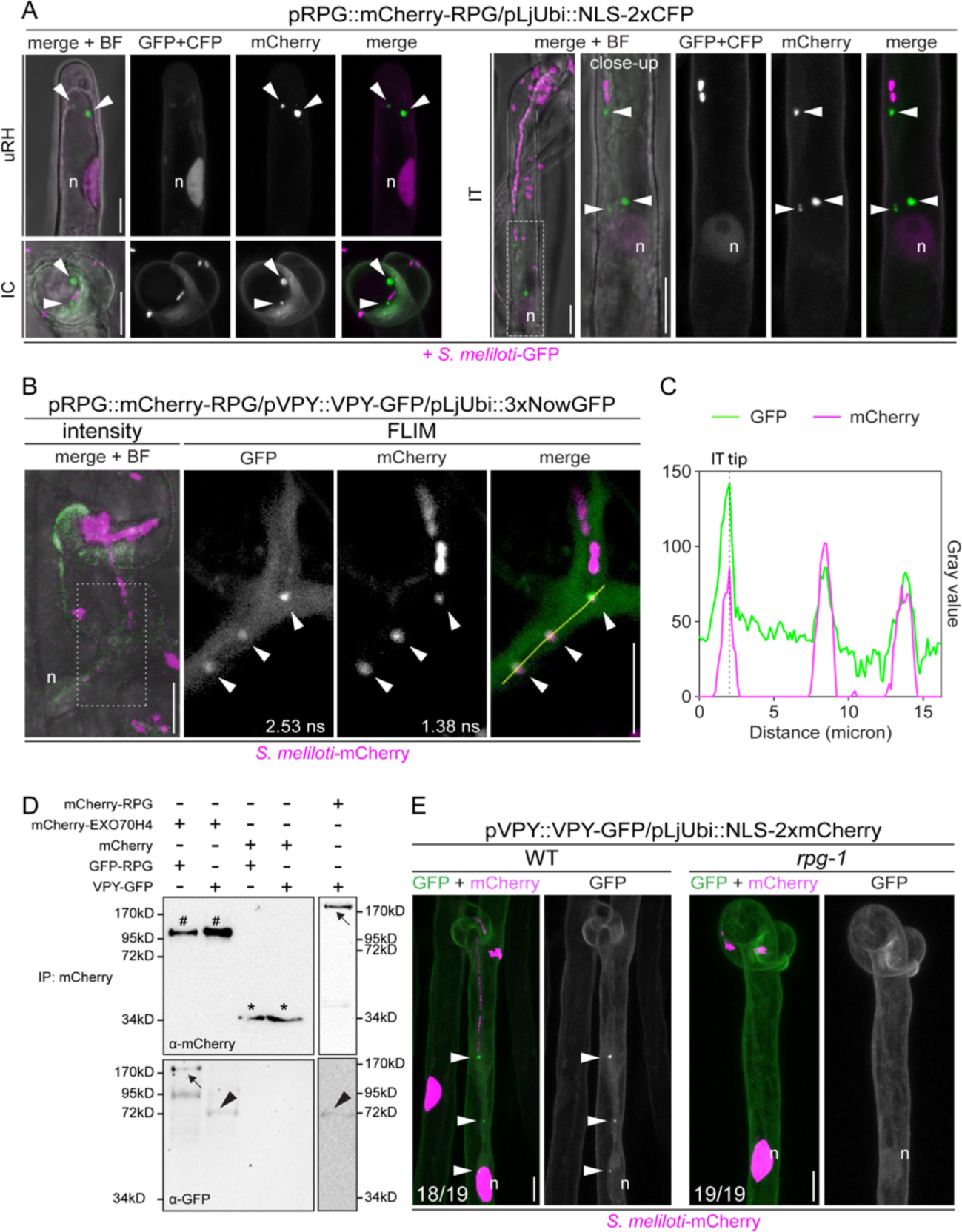
RPG is a key component of the VPY-LIN-EXO70H4 infectosome complex. **(A)** Live-cell confocal images showing mCherry-RPG localization in root hairs of WT transgenic roots at 4-5 dpi with *S. meliloti-*GFP. mCherry-RPG accumulates in cytoplasmic punctate structures (arrowheads) which are found close to the apex of uninfected root hairs (uRH), in the vicinity of rhizobia entrapped within the infection chamber of curled root hairs (IC) and associated to the perinuclear cytoplasm and the tip of infection threads (IT). The dashed white line box indicates the region shown in the close-up. A nuclear-localized tandem CFP was used as transformation marker. Images are intensity projections. Individual channels are false colored in grey. When channels are merged, the GFP/CFP channel is shown in magenta and the mCherry channel is shown in green. The bright field (BF) is overlaid with merged fluorescent channels. **(B)** Co- localization of mCherry-RPG with VPY-GFP in cytoplasmic and IT tip-associated puncta (arrowheads) in root hairs of WT transgenic roots imaged at 4-5 dpi with *S. meliloti*-mCherry. The intensity-based image is a merge of maximum intensity projections of GFP (green) and mCherry (magenta) channels overlaid with the brightfield (BF) channel. The dashed white line box indicates the region shown in FLIM images. FLIM images are maximum intensity projections of two focal planes. Lifetime values (ns) obtained from exponential reconvolution of decay profiles of GFP and mCherry channels are reported. Individual channels are shown in grey. When components are merged, GFP is shown in green and mCherry in magenta. A nuclear-localized triple NowGFP was used as transformation marker (not visible in these images). **(C)** Intensity profile of GFP and mCherry signal along the yellow transect shown in **(B)**. The intensity peaks at the infection thread tip (IT tip) are marked by a dashed black line **(D)** Co-immunoprecipitation analysis using anti-RFP nanotraps showing interaction between GFP-RPG and mCherry-EXO70H4 and between mCherry-RPG and VPY-GFP. Total proteins were extracted from protoplasts obtained from *N. benthamiana* leaves co-expressing RPG, VPY and EXO70H4 in different combination of pairs. Black arrows indicate mCherry-RPG (upper right panel) and GFP-RPG (lower left panel), hashtags indicate mCherry-EXO70H4, arrowheads indicate VPY, asterisks indicate mCherry. **(E)** Localization pattern of VPY-GFP in root hairs of WT and *rpg-1* transgenic roots imaged at 4-7 dpi with *S. meliloti*- mCherry. VPY-positive puncta (arrowheads) are systematically present in WT but not in *rpg-1* infected root hairs. Numbers indicate frequencies of observations, made on a total number of 8 (WT) and 10 (*rpg-1*) composite plants. A nuclear-localized tandem mCherry was used as transformation marker. GFP channel is shown in grey when isolated, in green when merged with mCherry (magenta). n = nucleus. Scale bars = 10 µm.

To evaluate the contribution of RPG protein domains to its punctate localization, we analyzed the subcellular patterning of the RPG deletion derivatives expressed under the control of the native *RPG* promoter in roots of composite *rpg-1 M. truncatula* plants. When expressed in this mutant background, mCherry-RPG-FL exhibited a comparable localization pattern to what was observed in the WT background (Figure 3-figure supplement 3A). We found that both mCherry-RPG-CC and mCherry-RPG-tCC accumulated to bright punctate structures as mCherry-RPG-FL, while the mCherry-RPG-NT signal remained diffuse in the cytoplasm and preferentially accumulated in the nucleus of transformed root hairs (Figure 3-figure supplement 3B-D). These results indicate that the coiled-coil domain is necessary and sufficient for RPG localization to discrete puncta and that deletion of residues from 331 to 473, although abolishing its functionality, does not affect its localization to these foci (Figure 2A-C and Figure 3-figure supplement 3B-C). In addition, the consistent nuclear accumulation of mCherry-RPG-NT compared to the occasional localization in this compartment that we observed for RPG-FL, suggests that the coiled-coil domain determines the preferential retention of the protein to cytoplasmic puncta (Figure 3-figure supplement 1C-D and Figure 3-figure supplement 3D).

This subcellular patterning resembles the recently described localization of the so-called “infectosome” complex formed by VPY, LIN and EXO70H4. These proteins systematically co- localize and interact in IT tip- and perinuclear-associated cytoplasmic puncta during root hair infection (Liu et al., 2019b; Roy et al., 2020). Thus, we tested if RPG is a component of this protein complex. For this, we first co-expressed mCherry-RPG and VPY-GFP under the control of their respective native promoters in roots of composite *M. truncatula* WT plants and imaged them at 4-5 dpi with *S. meliloti*-mCherry. Since the signal from co-expressed RPG- and VPY- fluorophore fusions were weak, to increase its detection and to unambiguously distinguish it from background fluorescence, we applied Fluorescent Lifetime Imaging Microscopy (FLIM). By performing multiple cycles of excitation and detection of photon arrival times, this method increases the intensity of fluorescence emission. Furthermore, it allows to analyze its decay profile, thereby enabling an increased spatial resolution of genetically encoded fluorophores according to their peculiar lifetimes (GFP ∼ 2.5-2.7 ns; mCherry ∼ 1.4-1.5 ns, Sarkisyan et al., 2015; Seefeldt et al., 2008) and the subtraction of the autofluorescence signal, having similar spectral characteristics but significantly shorter lifetimes (see materials and methods for more details). We found that mCherry-RPG clearly co-localized with VPY-GFP in punctate structures located at the tip of the IT and in the cytoplasm connecting the nucleus to the IT tip of infected root hairs (Figure 3B-C). A VPY-GFP signal of weaker intensity was also diffusely visible in the cytoplasm, as previously reported (Liu et al., 2019b) (Figure 3B-C). Co-expression of GFP-RPG with mCherry-LIN, the latter under the control of the constitutive *Ubiquitin* promoter from *L. japonicus* (pLjUbi), also resulted in distinct co-localization of the two proteins in root hairs undergoing infection (Figure 3-figure supplement 4A-B). Here, FLIM enabled a clear separation of mCherry-LIN from the signal of the NLS-3xmScarlet transformation marker (lifetime ∼ 3.5-3.9 ns, Bindels et al., 2016, 2020) (Figure 3-figure supplement 4A-B).

Furthermore, we performed co-immunoprecipitation (Co-IP) assays using protoplasts obtained from agroinfiltrated *N. benthamiana* leaves constitutively co-expressing RPG, VPY and EXO70H4 in different combinations of pairs. Using anti-mCherry nanobody traps on protein extracts from protoplasts, revealed that GFP-RPG and mCherry-RPG co-purified with mCherry-EXO70H4 and VPY-GFP, respectively (Figure 3D), indicating that these proteins indeed belong to the same complex.

Since the characteristic punctate localization of VPY is triggered by rhizobial inoculation and does not depend on functional LIN (Liu et al., 2019a), we hypothesized that VPY accumulation in these puncta may be triggered by and dependent on RPG, as *RPG* expression is induced in root hairs starting from 2 days following inoculation with *S. meliloti* (Arrighi et al., 2008; Liu et al., 2019a). To test this, we expressed VPY-GFP in roots of WT and *rpg-1* composite plants and visualized its spatial patterning in root hairs at 4-7 dpi with *S. meliloti*-mCherry. Indeed, while we systematically observed bright VPY-positive puncta in infected (18/19 in a total number of 8 plants) and uninfected (6/8 in a total number of 6 plants) root hairs of transformed WT roots, we never detected such punctate subcellular structures in *rpg-1* transformed root hairs, neither infected (19/19 events in a total number of 10 plants) nor uninfected (8/8 events in a total number of 7 plants). In all cases VPY exhibited a diffuse cytoplasmic signal (Figure 3E and Figure 3-figure supplement 5A). In few instances (5/19 infection events found in 2 plants out of 10), we observed VPY accumulating close to bacteria entrapped within a curl, seemingly surrounding the site of membrane invagination occurring at the early stage of IT emergence, but no cytoplasmic or perinuclear-associated puncta were visible (Figure 3-figure supplement 5B). This is in sharp contrast to what we observed in WT root hairs at a similar stage of infection (Figure 3-figure supplement 5B).

From these results we conclude that RPG is a key component of the infectosome machinery controlling IT polar growth, being necessary to modulate rhizobial-induced focusing of VPY to IT- tip and perinuclear-associated puncta during this process. The predicted function of the infectosome and the loss of VPY recruitment to IT-tip associated puncta in *rpg-1,* which fails to develop thin and elongated infection threads, strongly suggest that RPG is required to sustain localized polarity at this specific symbiotic domain.

### RPG is required to maintain PI(4,5)P2 enrichment on the membrane of the IT tip

In tip-growing plant cells such as root hairs and pollen tubes, confined polarity at the apical membrane domain is sustained by local enrichment of the low abundant phosphoinositide species PI(4,5)P2, which serve as landmarks to recruit diverse protein effectors regulating cytoskeleton rearrangements, vesicle trafficking and signaling (Ischebeck et al., 2010; Noack & Jaillais, 2020). As the polarized IT tip likely recruits part of the machinery controlling polarity establishment during root hair growth (Gage, 2004), we hypothesized that local enrichment of PI(4,5)P2 occurs at the site of symbiont accommodation. To test this, we expressed the well-characterized genetically encoded PI(4,5)P2 biosensor mCitrine-2xPH^PLC^ (Simon et al., 2014) under the control of the infection-induced *ENOD11* promoter (Boisson-Dernier et al., 2005; Journet et al., 2001) in roots of WT *M. truncatula* composite plants and analyzed its localization patterns following inoculation with *S. meliloti*-mCherry (4-8 dpi). To increase the robustness of our analysis, we adopted a systematic FLIM-based approach that allowed us to i) unambiguously identify local membrane enrichment of the mCitrine-PH^PLC^ probe (Lifetime ∼ 3 ns; Söhnel et al., 2016) and discriminate it from cell wall autofluorescence (Lifetime ≤ 1.1 ns) and ii) integrate lifetime measurements obtained from the observation of multiple events in a visually-intuitive form using the Phasor approach (Malacrida et al., 2021; Ranjit et al., 2018) and statistically compare them (Figure 4-figure supplement 1A; see materials and methods for full description).

**Figure 4.**
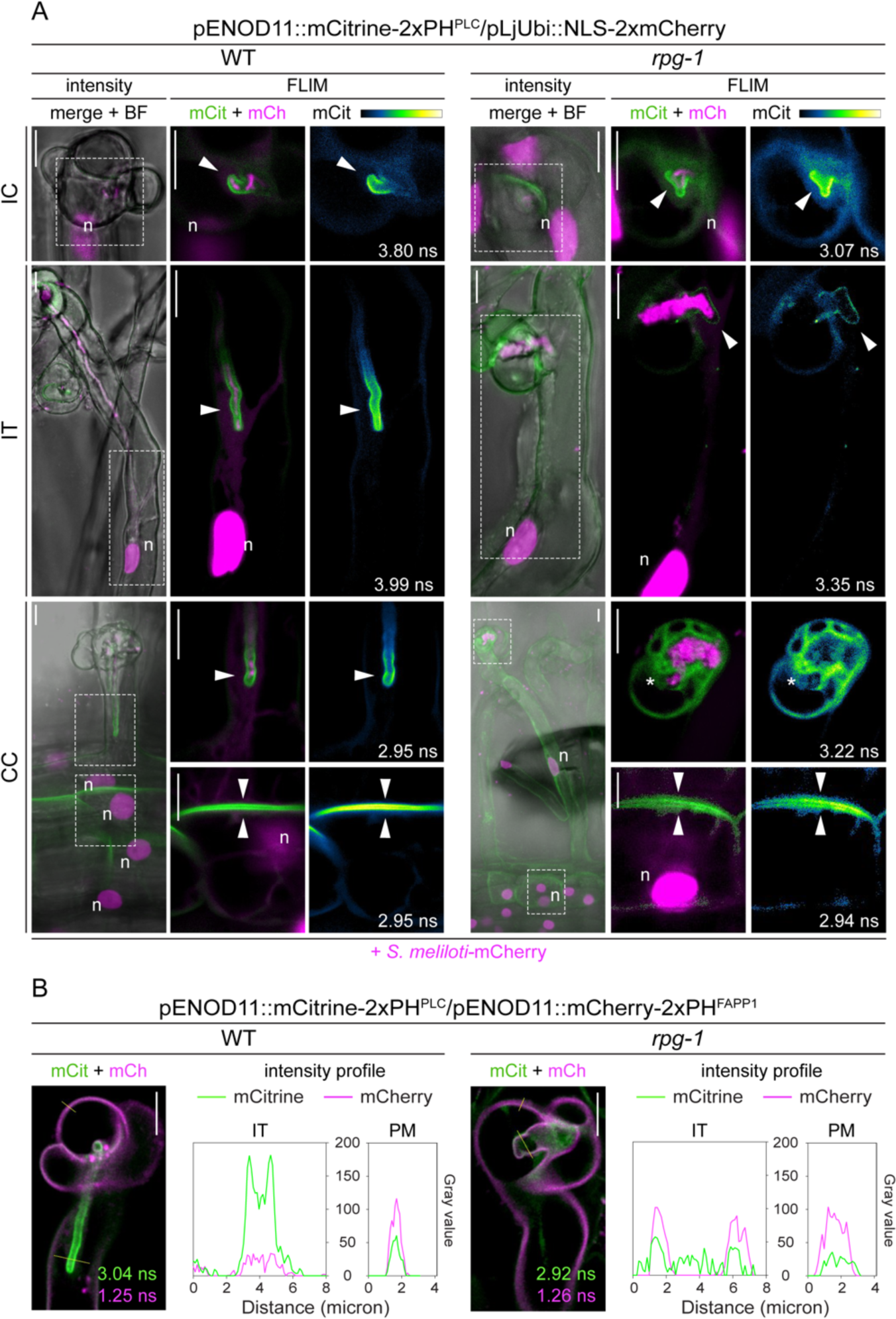
RPG is required to maintain strong membrane polarization at the IT tip. **(A)** Live-cell confocal images of root hairs from transgenic WT and *rpg-1* roots expressing the PI(4,5)P2 biosensor mCitrine-2xPH^PLC^ at 4-8 dpi with *S. meliloti*-mCherry. The distribution of mCitrine-2xPH^PLC^ in WT and *rpg-1* transgenic roots is compared during emergence of rhizobia from the infection chamber (IC), in root hairs hosting infection threads (IT) and in cortical cells underlying epidermal infection events (CC). Membrane domains where differential enrichment of PI(4,5)P2 occurred are indicated with arrowheads. Cytoplasmic signal of the biosensor in *rpg-1* infected root hairs is indicated with an asterisk. Intensity- based images are a merge of intensity projections of mCitrine (green) and mCherry (magenta) channels overlaid with the brightfield (BF) channel. The dashed white line box in intensity images indicates the region shown in FLIM images. FLIM images are single focal planes; the mCitrine component is shown in green when merged with the mCherry component (magenta) and in Green Fire Blue when isolated, with yellow indicating the maximum intensity and blue a low level of fluorescence. Lifetime values (ns) obtained from exponential reconvolution of decay profiles of the mCitrine channel are reported. A nuclear-localized tandem mCherry was used as transformation marker. n = nucleus. Scale bars = 10µm. **(B)** Co-visualization of PI4P and PI(4,5)P2 in root hairs harboring ITs from WT and *rpg-1* transgenic plants co- expressing mCitrine-2xPH^PLC^ and mCherry-2xPH^FAPP1^ at 4 dpi with *S. meliloti*-LacZ. Images are merge of mCitrine (green) and mCherry (magenta) components with indicated lifetime obtained from exponential reconvolution of decay profiles of the respective channels. Intensity profiles show grey values along the yellow transects intersecting the infection thread membrane (IT) and the root hair plasma membrane (PM) in the corresponding images. Scale bars = 10 µm.

In uninfected growing root hairs, PI(4,5)P2 enriched at the apical membrane domain (Figure 4-figure supplement 2A and Figure 4-figure supplement 4A) while at the onset of infection PI(4,5)P2 was locally and specifically enriched at the membrane surrounding invading bacteria emerging from the infection chamber (IC) (Figure 4A – IC and Figure 4-figure supplement 4B). This indicates that symbiont accommodation coincides with membrane polarization at this host-microbe interface. In root hairs harboring actively growing ITs, we consistently observed local enrichment of the phosphoinositide on the membrane surrounding the tip and the terminal part of the extending tube, connected to the nucleus by a dense column of cytoplasm (Figure 4A – IT and Figure 4-figure supplement 4C), confirming that bacterial progression is associated with strong polarization of the IT membrane. Notably, we observed that local PI(4,5)P2 enrichment also occurred on the basal membrane of root hairs hosting polarized ITs extending within a cytoplasmic column connected to the root hair base, and on the apical membrane of underlying cortical neighboring cells (Figure 4A – CC, Figure 4-figure supplement 2B and Figure supplement 4D). This fully coincided with the reorganization of the cortical cell cytoplasm into a dense central column occupied by the nucleus, which is typical of pre-infection thread (PIT) responses (Van Brussel et al., 1992) (Figure 4A - CC and Figure 4-figure supplement 2B). This demonstrates that local membrane polarization occurs on the trajectory of IT propagation and might serve to direct local changes of the cell wall and apoplastic space in preparation for the transcellular passage of the bacteria.

To further verify that the observed membrane localization of the 2xPH^PLC^ biosensor reflects PI(4,5)P2 enrichment, we generated a mutant version of the probe bearing amino acid substitutions R37D and R40D, which substantially reduce its PI(4,5)P2-binding capacity and abolish its interaction with the plasma membrane (mCitrine-2xPH^PLC-mut^; Yagisawa et al., 1998; Ivanov & Harrison, 2019). Expression of this mutated probe under the control of the *ENOD11* promoter in WT transgenic roots resulted in exclusively diffuse cytoplasmic signal in both uninfected root hairs (Figure 4-figure supplement 3 - uRH), infected root hairs (Figure 4-figure supplement 3 – IC and IT) and in cortical cells underlying primary infection events (Figure 4-figure supplement 3 - CC) imaged at 4-7 dpi with *S. meliloti*-mCherry.

Next, we evaluated the impact of a loss of RPG on infection-dependent membrane repolarization by expressing the PI(4,5)P2 probe in *rpg-1* transgenic roots and monitoring its distribution pattern upon inoculation (*S. meliloti* mCherry, 4-8 dpi). Here, the apex of uninfected growing root hairs was labelled by the PI(4,5)P2 biosensor as in the WT (Figure 4-figure supplement 2A and Figure 4-figure supplement 4A). PI(4,5)P2 patterning was also similar to WT at the very initial stage of bacterial accommodation, where we observed a local enrichment occurring on the membrane surrounding the bacteria protruding from the IC (Figure 4A – IC and figure 4-figure supplement 4B). Thus, RPG is not required to establish membrane polarization at this stage. However, PI(4,5)P2 failed to strongly enrich on the majority of membranes (21/33 IT from 10 plants) of typically enlarged infection threads developed by the *rpg-1* mutant. Here, the signal of the probe remained either low and uniformly labelling the membrane, cytoplasmic or almost absent (Figure 4A, Figure 4-figure supplement 2C and Figure 4-figure supplement 4C), indicating that functional RPG is required to maintain IT polarization during symbiont accommodation. When present, the enrichment was often found on short membrane protrusions formed at multiple polar sites on the enlarged infection structures developed within *rpg-1* root hairs, further supporting that not the establishment but rather the maintenance and restriction of symbiont-induced polarization is RPG-dependent (Figure 4-figure supplement 2C and Figure 4-figure supplement 4C). In addition, we found that development of polar domains at the base of infected root hairs and on the apical membrane of underlying cortical cells was not affected in *rpg-1* (Figure 4A – CC and Figure 4-figure supplement 4D), implying that RPG is not necessary for these pre-infection responses but exclusively for IT maintenance and progression. This is in agreement with previous findings showing that PIT formation and microtubule rearrangements in cortical cells of *rpg-1* are not perturbed (Arrighi et al., 2008). However, such responses were uncoupled from IT polarization and nuclear-guided progression of the tube within the root hair (Figure 4A-CC), suggesting that functional RPG is important to maintain synchronization between cells during the infection process.

To provide statistical support to our analysis, we compared the Phasor fingerprint of fluorescence emission (see Materials and Methods for a detailed description) at different membrane domains analyzed in multiple cells of different plants for each genotype (Figure 4-figure supplement 4E). A statistical difference was only found on the Phasor position of fluorescence emission at the IT membrane but not at other domains (Figure 4-figure supplement 4E).

To further characterize and compare the membrane identity of WT and *rpg-1* ITs, we co-visualized the distribution of PI(4,5)P2 together with its synthetic precursor PI4P, which is highly enriched at the plasma membrane (PM) of plant cells and contributes to define its identity (Simon et al., 2016). For this, we co-expressed mCitrine-2xPH^PLC^ with the PI4P biosensor mCherry-2xPH^FAPP1^ (Simon et al., 2014) both under the control of the *ENOD11* promoter in roots of WT and *rpg-1* composite plants. To avoid that the intense signal from fluorescently-labelled bacterial symbionts could interfere with visualization of any of the two biosensors, we inoculated composite plants with a non-fluorescent rhizobial strain (*S. meliloti*-LacZ).

In WT infected root hairs, we found that the tip-polarized enrichment of the PI(4,5)P2 biosensor on the IT membrane spatially coincided with low enrichment of PI4P probe, which instead preferentially accumulated on the root hair PM (Figure 4B and Figure 4-figure supplement 5). Such differential membrane enrichment of the two biosensors was much reduced in *rpg-1* infected root hairs, where PI4P was similarly enriched on the root hair PM and on the infection thread membrane, the latter being characterized by low accumulation of the PI(4,5)P2 probe (Figure 4B and Figure 4-figure supplement 5). These observations support our previous set of data, clearly indicating a role for RPG in sustaining localized membrane polarization at the tip of the IT.

Since PI(4,5)P2-enriched membrane domains constitute important platforms steering cytoskeleton rearrangements and targeted vesicle delivery (Bloch et al., 2016; Doumane et al., 2020; Stanislas et al., 2018; Synek et al., 2021) and as a last set of experiments to verify the role of RPG in maintaining localized polarity at the IT tip, we decided to test how microtubule organization and polar secretion associated to this symbiotic domain are affected in *rpg-1*.

### Tip-to-nucleus microtubule connectivity is perturbed in *rpg-1*

During IT growth, a dense array of endoplasmic microtubules (EMTs) connects the IT tip to the nucleus migrating towards the root hair base and maintenance of such functional connection is thought to be necessary to guide IT progression within host cells (Timmers et al., 1999; Perrine- Walker et al., 2014; Fournier et al., 2008; Faehraeus, 1957).

Since nuclear movement appeared uncoupled from IT growth in *rpg-1* (Figure 4A - CC), we tested if the tip-to-nucleus connectivity is perturbed in *rpg-1* by comparing microtubule organization in WT and *rpg-1* root hairs harboring ITs using live-cell imaging. To reduce possible perturbations of microtubule dynamics given by ectopic expression of exogenous tubulin isoforms (Celler et al., 2016), we labelled microtubules expressing an N-terminal GFP fusion of an endogenous tubulin beta isoform (*Medtr8g107250,* hereafter SYMTUB) up-regulated in infected root hairs (Breakspear et al., 2014) under the control of a native 1.6kb promoter (pSYMTUB::eGFP-SYMTUB) in WT and *rpg-1* roots. In most of WT root hairs hosting growing ITs imaged at 4-8dpi with *S. meliloti*-mCherry, EMTs were organized in a dense cone-shape array anchored to the nucleus and extending towards the IT tip, keeping a tight connection between these two poles (Figure 5A). This patterning is consistent with previous reports (Timmers 1999; Perrine-Walker et al., 2014), supporting the validity of our marker. When imaging *rpg-1* infected root hairs at the same time points, such connection appeared loosened in the majority of cases, as EMTs associated to the IT mostly appeared dispersed and stretched, no longer bridging the tip and the nucleus and often positioned at a consistent distance away from the infection site (Figure 5A). Notably, the IT tip often appeared mis-oriented with respect to the direction of growth, pointing towards the root hair shank instead of following the repositioning of the nucleus down the root hair, suggesting that loss of nuclear-mediated IT guidance occurs in *rpg-1*. This indicates that RPG is required to maintain a functional physical connection between the nucleus and the extending tip.

**Figure 5.**
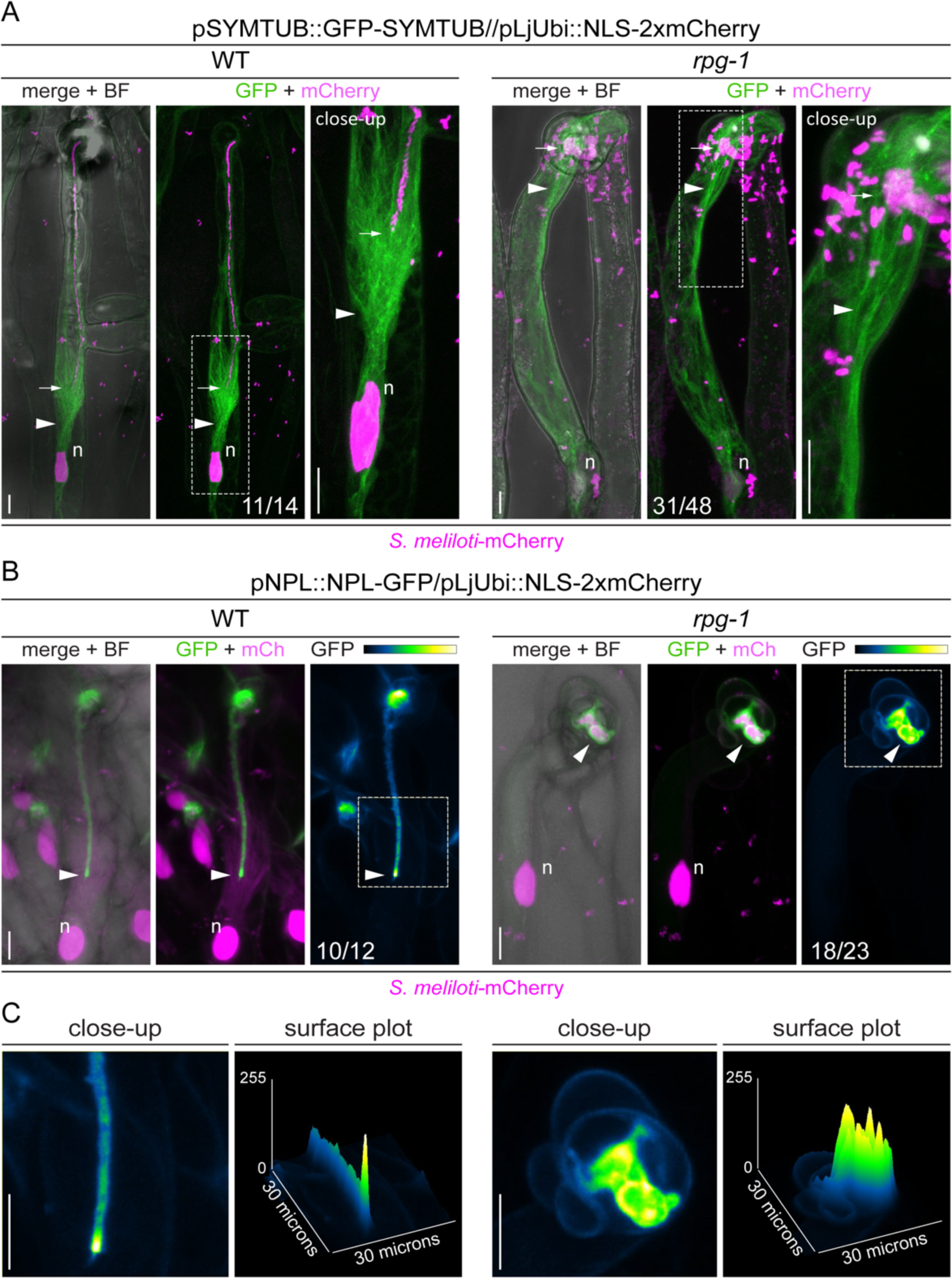
The tip-to-nucleus microtubule connectivity and polar secretion of NPL are affected in *rpg-1*. **(A)** In vivo imaging of microtubule patterning in root hairs hosting infection threads from WT and *rpg-1* transgenic roots expressing GFP-SYMTUB at 4-8 dpi with *S. meliloti*-mCherry. The endoplasmic microtubule array (arrowhead) bridging the nucleus (n) and the IT tip (arrow) in WT root hairs appears stretched in *rpg-1* root hairs, where the IT tip exhibit a tilted direction of growth. Images are merge of intensity projections of GFP (green) and mCherry (magenta) channels. The brightfield (BF) channel is overlaid with the merge. The dashed white line boxes indicate the region shown in the close-up. Numbers indicate frequencies of observation made on a total number of 6 (WT) and 14 (*rpg-1*) composite plants. **(B)** Live-cell confocal images showing focal accumulations of NPL-GFP at the IT tip (arrowhead) in WT root hairs in contrast to its unrestricted distribution in the apoplastic space surrounding ITs developed within root hairs of *rpg-1* transgenic roots. Images are intensity projections. The GFP channel is shown in green when merged with the mCherry channel (magenta), or in Green Fire Blue when isolated, with yellow indicating the maximum intensity and blue a low level of fluorescence. The brightfield (BF) channel is overlaid with the merge. Numbers indicate frequencies of observations made on a total number of 6 plants (WT and *rpg-1*). **(C)** Close-up and surface plot of the regions bounded by the dashed white line boxes in **(B).** The signal of NPL-GFP in WT ITs is described by a single peak compared to multiple peaks detected in *rpg-1.* Scale bars = 10 µm.

### Polar secretion of NPL requires functional RPG

The cell wall modifying enzyme NPL, which is essential for IT initiation and elongation in both *L. japonicus* and *M. truncatula* (Xie et al., 2012; Liu et al., 2019a) is secreted to the IT apoplast and accumulates particularly at IT tip regions in *M. truncatula* (Liu et al., 2019a), strongly suggesting that its tip-targeted secretion is required for IT polar extension. To evaluate the role of RPG in NPL targeting, we compared the localization pattern of this enzyme in WT and *rpg-1* infected root hairs. For this, we expressed an NPL-GFP fusion under the control of the native *NPL* promoter in WT and *rpg-1* roots and imaged infected root hairs at 4-8 dpi. In WT root hairs harboring growing ITs and in agreement with a previous report (Liu et al., 2019a), NPL consistently showed a preferential accumulation in the apoplastic space surrounding the tip when compared to the older parts of the IT (Figure 5B-C). By contrast, NPL strongly accumulated in the whole lumen of the enlarged ITs without any increased accumulation at a specific site in most *rpg-1* root hairs (Figure 5B-C). This indicates that RPG is required to restrict the secretion domain of NPL during infection.

## DISCUSSION

The capacity to intracellularly accommodate nitrogen-fixing bacteria is a hallmark of the RNS. Bacterial internalization requires profound restructuring of the host cell to form an infection thread compartment via polarized secretion of plasma membrane and cell wall material to a single focal point. Polar progression of IT structures determines the establishment of a functional nitrogen-fixing association.

Here, we demonstrate that RPG is a crucial component of the infectosome machinery sustaining polar growth of intracellular ITs (Figure 3A-C and Figure 3-figure supplement 4 A-B), being necessary to recruit VPY to the IT-tip and perinuclear puncta during IT progression within root hairs (Figure 3E and Figure 3-figure supplement 5A-B). A cooperative function of RPG with the infectosome complex in regulating IT polar growth is consistent with the phenotypes of *rpg*, *vpy* and *lin* mutants, exhibiting strong defects in the establishment and/or the maintenance of IT progression but not being affected in nodule organogenesis (Kiss et al., 2009; Murray et al., 2011; Liu et al., 2019a, Figure 1B-D and Figure 1-figure supplement 1A-B). Moreover, the peculiar enlargement of ITs developed by *rpg* mutants, indicative of a loss of unidirectional polarized growth, is well explained by the failure of *rpg-1* to maintain the typical VPY punctate localization observed in WT root hairs (Figure 3E and Figure 3-figure supplement 5A-B). Interestingly, infectosome-like and VPY-positive compartments have also been observed during the arbuscular mycorrhiza symbiosis (Pumplin et al., 2010; Zhang et al., 2015) and *vpy* mutants exhibit defects in this association (Feddermann et al., 2010; Pumplin et al., 2010). Given the fact that VPY-recruitment into infectosome structures is abolished in an *rpg* mutant, it is tempting to speculate that RPG serves as a specificity determinant for infectosome- related endo- and exocytotic events during RNS, which is supported by the transcriptional dependency of *RPG* from NIN (Liu et al., 2019a; Soyano & Hayashi, 2014) and is in line with *RPG* being one of the few essential factors that have been lost in non-nodulating species of the FaFaCuRO clade (Griesmann et al., 2018; van Velzen et al., 2018). Upon recruitment of VPY into the infectosome complex it is further stabilized by LIN (Liu et al., 2019b) and this can also be assumed for CERBERUS, the ortholog of LIN in *L. japonicus*, which has been shown to stabilize VPY protein levels (Liu et al., 2021). Different to *RPG*, the genes encoding for VPY and LIN are retained in angiosperm species forming intracellular symbioses (Radhakrishnan et al., 2020) and at least *VPY* is additionally required for intracellular accommodation of the fungal partner during arbuscular mycorrhizal symbiosis (Feddermann et al., 2010; Pumplin et al., 2010).

While the existence of infectosomes is supported by this and other studies (Liu et al., 2019b; Zhang et al., 2015), the full nature of this presumably membrane-less compartment remains to be investigated further. It is, however, tempting to draw parallels to the so called “polarisome”, a macromolecular complex regulating polarized growth of yeasts and fungal hyphae (Xie & Miao, 2021). Polarisomes are dense spot-like structures localized at the site of polarized growth and are formed by a number of core components and many cooperative interactors (Xie & Miao, 2021). The polarisome is also required to maintain the Spitzenkörper (Crampin et al., 2005), a vesicle supply center located at the tip during hyphal growth, whose positioning is correlated with growth directionality (Riquelme & Sánchez-León, 2014). Interestingly, a myosin protein has recently been shown to mediate focal packing of the polarisome (Dünkler et al., 2021). The structural similarity of the RPG C-terminal extension with myosins, as supported by structural predictions and as previously suggested (Arrighi et al., 2008), and the inability of VPY to localize into distinct infectosome foci, further supports the hypothesis of RPG functioning as an organizational hub protein for infectosome assembly and polarity maintenance during intracellular rhizobial infections. It should, however, be considered that a vesicle-associated fraction of RPG as well as of the other infectosome components outside the puncta might contribute to their functionalities, although these structures cannot be visualized at present due to their size and/or fluorescence intensity. A possible association of RPG with vesicular compartments would be compatible with the predicted membrane-binding capacity of its NT-C2 domain (D. Zhang & Aravind, 2010).

Rendering the infectosome complex dysfunctional in *rpg* mutants results in the loss of the highly polarized membrane enrichment of PI(4,5)P2 (Figure 4A-B) and most likely in mistargeted secretion of key factors such as NPL required for IT progression to the growing IT tip (Liu et al., 2019a; Xie et al., 2012). It is noteworthy that maintenance of the IC and secretion of NPL into this apoplastic compartment is unaltered in *rpg* mutants, but is unconfined during IT development (Figure 5B-C). As in many eukaryotic polarized systems, where localized exocytosis targets PI(4,5)P2-rich membrane domains, accumulation of this phosphoinositide was also shown to occur on perimicrobial membranes during pathogenic infections (Qin et al., 2020) and during AM symbiosis, where PI(4,5)P2 specifically accumulated at the tips of intracellular linear hyphae (Ivanov & Harrison, 2019). Taken together, our data strongly support the hypothesis that IT polarity requires few specific factors like RPG but predominantly relies on a highly conserved mechanism as suggested earlier (Gage, 2004; Yamazaki & Hayashi, 2015). While the secretion domain in curled root hairs comprises the entire IC (Fournier et al., 2015), the loss of highly restricted focal polarization results in the formation of enlarged ITs (Figure 1B-D). Most of these ITs progress in an undirected manner through the root hairs of *rpg* mutants, as indicated by the loss of microtubule-mediated connectivity between the nucleus and the IT tip (Figure 5A). Although polarization of the basal membrane domain in the respective trichoblasts and formation of PITs in the outer cortical cells are unaltered (Figure 4, (Arrighi et al., 2008), effective infection of nodules is impaired (Figure 1-figure supplement 1B) likely because a strictly defined time window that allows transcellular IT growth through recently divided cells is missed by these slowly growing ITs in the *rpg* mutant.

## MATERIALS AND METHODS

### Plant material and phenotypic analysis

*M. truncatula* wild-type ecotypes R108, Jemalong A17 and two *rpg* mutant alleles were used in this study. *rpg-1* is a previously described EMS mutant bearing a premature stop codon in the third exon of the RPG gene (Arrighi et al., 2008). Seeds of this allele were kindly provided by Clare Gough (Institut National de la Recherche Agronomique de Toulouse, Toulouse, France). *rpg-2* is a *Tnt-1* insertion line (NF11990) and was obtained from the Noble Research Institute (OK, USA).

Seeds were scarified by soaking them in sulfuric acid (H2SO4) 96% for 8 minutes, then rinsed six times with sterile tap water and surface sterilized with a solution of 1.2% sodium hypochlorite (NaClO) and 0.1% sodium dodecyl sulfate (SDS) for 1 minute. After rinsing six times with sterile tap water, seeds were transferred to 1% agar/water plates and vernalized at 4° in dark for 4-5 days. Seeds were then moved to 24°C in dark for 10hrs to induce germination.

For phenotypical analysis of IT morphology, germinated seedlings were transferred to pots containing a mixture (1:1 v/v) of quartz sand (grain size 0.1-0.5 mm, Sakret) and vermiculite (0-3 mm, Terra Exotica), inoculated with *S. meliloti* (Sm2011, OD600=0.001, 5 mL per pot) and grown in a controlled environment chamber at 24°C with a 16/8 hours light/dark photoperiod, and 70% humidity. Pots were watered twice a week with tap water and fertilized once a week with a ¼ Hoagland solution containing 0.1mM KNH3. Plants were harvested at 35 dpi, the root system was excised and fixed in 4% PFA (paraformaldehyde). Prior to imaging, roots were washed three times with 1 x PBS (phosphate-saline buffer). Infected root hairs were imaged using a confocal microscope and measurement of the infection thread diameter was performed with ImageJ/(Fiji) software (Schindelin et al., 2012). Infection thread diameter has been scored on 19-31 root hairs of at least 4 individual plants.

For morphological examination of cortical infection threads, germinated seedlings were transferred to sterile plates containing solid Fahräeus medium supplemented with 0.1 mM NH4NO3 and grown vertically in a controlled environment chamber at 24°C with a 16/8 hours light/dark photoperiod for 1 week before inoculating them with *S. meliloti* (RCR2011, OD600=0.01). At 14 dpi the plant root system was excised, fixed in 4% PFA under vacuum for 15 min and then washed twice with 1x PBS. Afterwards, roots were covered with ClearSee solution (Ursache et al., 2017) and incubated for 7-10 days. Prior to imaging, roots were stained with Calcofluor white for 45 minutes, washed twice with ClearSee and incubated for 1hr in the last wash. Cortical infection threads were then imaged using a confocal microscope.

For phenotypic analysis of nodulation capacity, germinated seedlings were transferred to pots containing Zeolite substrate (50% fraction 1.0-2.5mm, 50% fraction 0.5-1.0-mm, Symbiom), inoculated with *S. meliloti* (RCR2011 pXLGD4, GMI6526, OD600=0.02, 10 ml per pot) and grown in a controlled environment chamber at 22°C with a 16/8 hours light/dark photoperiod, and 70% humidity. Pots were watered once a week with tap water and fertilized once a week with Fahräeus medium (0.5 mM NH4NO3). Plants were harvested 21dpi. Two independent replica were performed.

### Hairy root transformation and rhizobial inoculation

Composite *M. truncatula* plants were obtained following the previously described procedure (Boisson-Dernier et al., 2001). In brief, transgenic *A. rhizogenes* (ARqua1) carrying the plasmid of interest was grown in liquid LB medium (5 mL) supplemented with appropriate antibiotic for 24hrs until reaching an OD600 of approximately 0.5-0.7. 300 μl of this culture were then spread on a plate containing solid LB medium with antibiotics and incubated at 28°C in dark for 48 hrs. The root meristem of germinated seedlings was removed with a scalpel and the wounded part was dragged on the *A. rhizogenes* solid culture before transferring seedlings to plates containing solid Fahräeus medium supplemented with 0.5 mM NH4NO3. Transformed seedling were grown vertically in a controlled environment chamber at 22°C in dark for 3 days and for additional 4 days with a 16/8 hours light/dark photoperiod providing shading to the root system. Afterwards, composite plants were transferred to new solid Fahräeus plates (0.5 mM NH4NO3) and grown for 10 days at 24°C with a 16/8 hours light/dark photoperiod. Transformed roots expressing the fluorescent selection marker were selected using a stereo microscope, untransformed roots were excised with a scalpel and composite plants transferred to either solid Fahräeus medium supplemented with 0.1 mM NH4NO3 (live-cell imaging of root hairs) or to pots (complementation assays).

*Sinorhizobium meliloti* (Sm2011) was grown in liquid TY medium (5 mL) supplemented with appropriate antibiotics for 3 days at 28°C. 100 μl of this culture were used as inoculum for a fresh culture (5mL), which was grown for 24 h at 28°C. A bacterial pellet was then obtained by centrifugation at 3000 rpm for 10 minutes, washed once with liquid Fahräeus medium (0.1 mM NH4NO3) and resuspended in the same medium to the reach a final OD600 = 0.01 (inoculation of composite plants in plates, 200 μl per plant ) or OD600 = 0.001 (inoculation in pots, 5 mL per plant).

### Construct design

Constructs used in this study were designed and assembled using Golden Gate cloning (Binder et al., 2014; Weber et al., 2011). All promoters, coding sequences and genomic sequences used were synthesized by Life Technologies. When internal BsaI or BpiI sites were present, individual point mutations were introduced *in silico*; in case of coding and genomic sequences, silent mutations were produced. A list of all L1 and L2 constructs including module composition, fluorophore linkers and tagging site is provided as supplementary file (Table S1).

For RPG, a sequence 1777 bp upstream of the start codon was used as promoter, as described earlier (Arrighi et al., 2008). The RPG coding sequence was used for complementation assays, subcellular localization studies and co-immunoprecipitation assays. RPG fragments corresponding to different deletion derivatives were synthesized by Life Technologies.

For VPY and EXO70H4, constructs were designed as described earlier (Liu et al., 2019b). For VPY, the genomic sequence and a 2926 bp promoter were used. The potato ST-LS1 intron-containing version of S65TGFP used as C-terminal tag to avoid leaking expression of the fluorophore in agrobacteria (Liu et al., 2019b) was synthesized by Life Technologies. For EXO70H4, the coding sequence was used.

The previously described pLjUbi::mCherry-LIN construct (Liu et al., 2019b) was used for co- localization studies.

For subcellular localization pattern of NPL, the coding sequence and a 2038 bp promoter were used. For microtubule fluorescent labelling, the coding sequence of SYMTUB was used and a sequence 1555 bp upstream of the start codon as promoter.

For expression of the phosphoinositide biosensors, a 1106 bp promoter of *ENOD11* was used. Sequences of 2xPH^PLC^ and 2xPH^FAPP1^, and of 2xPH^PLC^ ^mut^ were designed according to (Simon et al., 2014) and (Yagisawa et al., 1998), respectively, and synthesized by Life Technologies.

The sequence data from RPG, VPY, EXO70H4, LIN, NPL, SYMTUB can be found in phytozome (https://phytozome.jgi.doe.gov/) with the following gene ID: RPG (Medtr1g090807), VPY (Medtr6g027840), EXO70H4 (Medtr4g062330), LIN (Medtr1g090320), NPL (Medtr3g086320),

SYMTUB (Medtr8g107250). The sequences of 2xPH^PLC^ and 2xPH^FAPP1^ biosensors can be downloaded at http://www.ens-lyon.fr/RDP/SiCE/PIPline.html.

### Complementation assays

Composite *M. truncatula* plants expressing the constructs of interest were transferred to pots containing a mixture of quartz sand and vermiculite (1:1 v/v), inoculated with *S. meliloti* (Sm2011, OD600=0.001, 5 mL per pot) and grown in a controlled environment chamber at 24°C with a 16/8 hours light/dark photoperiod, and 70% humidity. Pots were watered twice a week with tap water and fertilized once a week with a ¼ Hoagland solution containing 0.1mM KNH3. Plants were harvested at 21dpi. Transformed roots exhibiting strong expression of the fluorescent marker were selected, excised and fixed in 4% PFA (paraformaldehyde) under vacuum for 15 min. After washing the roots twice with 1x PBS (phosphate-saline buffer), roots were covered with ClearSee solution (Ursache et al., 2017) and incubated for 7-10 days. ClearSee solution was refreshed every two days. Prior to imaging, roots were stained with Calcofluor white for 45 minutes, washed twice with ClearSee and incubated for 1h in the last wash. Infected root hairs were imaged using a confocal microscope and measurement of the infection thread diameter was performed with ImageJ/(Fiji) software (Schindelin et al., 2012). Infection thread diameter has been scored on 20-32 independently transformed roots from at least 5 plants.

### Live-cell imaging of root hairs

Composite plants growing on solid Fahräeus medium supplemented with 0.1 mM NH4NO3 were inoculated with *S. meliloti* (Sm2011) by adding 200 μl of bacterial suspension (OD600 = 0.01) on each root system. Prior to inoculation, the position of the apex of transformed roots was marked on plates. 4-8 dpi, root segments that had grown below the mark were excised, mounted in water and root hairs were imaged with a confocal microscope. For live-cell imaging experiments, at least two independent replica were conducted with six composite plants (arising from independent transformation events) analyzed per condition, unless otherwise stated.

### Confocal Laser-Scanning Microscopy (CLSM) and Fluorescence Lifetime Imaging Microscopy (FLIM)

All imaging was performed using a Leica SP8 FALCON confocal microscope equipped with a 20x or 40x water immersion lens (Leica Microsystems, Mannheim, Germany). A White Light Laser (WLL) was used as excitation source. GFP and mCitrine were excited at 488nm and the emission collected at 500-550nm. mCherry and mScarlet were excited at 581nm and emission detected at 590- 640nm. To excite calcofluor white, a 405nm laser diode was used and emission was detected at 425- 475nm.

For Fluorescence Lifetime Imaging Microscopy, a Leica GaAsP-hybrid detector was used to collect fluorescent emission. Different channels were acquired in sequential scanning between frames. A pulse repetition rate of 80mHz was used with the exception that images in Figure 3B and Figure 3 – figure supplement 1A were acquired using a 40mHz pulse repetition rate for the GFP/NowGFP channel and the mCherry/mScarlet channel, respectively. Fluorescence emission was acquired from single focal planes using 20 scan repetitions for each channel.

### Intensity-based and FLIM-based image analysis

Intensity-based images were analyzed using ImageJ/(Fiji) software (Schindelin et al., 2012).

FLIM-based images were analyzed using LAS X SP8 Control Software (Leica Microsystems GmbH). Global fitting of the intensity decay profile using n-exponential reconvolution was performed to separate major fluorescent components within each channel and calculate their lifetime. The number of components (n) used for curve fitting was determined according to evaluation of the chi-squared (χ2) value (Lakowicz, 2006) and a threshold of 30 photons was applied to generate the final images. Components with lifetime value < 1.2 ns representing autofluorescent species accumulating in plant cell walls (Donaldson, 2020; Heskes et al., 2012) were subtracted from each channel.

All data on reconvolution of decay profiles of images presented in the article are provided as source files.

Since global fitting does not always allow to resolve and characterize the decay profile of pixels constituting minor populations within an image (Ranjit et al., 2018), the decay profile of fluorescent emission of regions of interest (ROIs) selected on membrane domains was additionally analyzed for the analysis and quantification of PI(4,5)P2 membrane enrichment. n-exponential fitting of the decay curve was carried out to resolve lifetime values associated to each ROI (Figure 4-figure supplement 1). Reconvolution data from all images and selected ROIs analyzed are provided as source files. Further, the raw decay profile of each ROI was analyzed using the Phasor approach to obtain a Phasor fingerprint whose position describes the fluorescent emission at that region and reflects its composition in terms of relative abundance of fluorescent species (Figure 4-figure supplement 1) (see Ranjit et al., 2018 and Malacrida et al., 2021 for detailed description of Phasor approach). Phasor plots were generated using a second harmonic, a threshold of 22 photons and a median of 15. To visualize and compare the Phasor fingerprint of ROIs selected in multiple images from different genotypes, the center of mass of Phasor images depicting the pixel populations originating from each ROI was calculated images using ImageJ/(Fiji) (Schindelin et al., 2012) and the obtained XM and YM coordinates were plotted on a graph (Figure 4-figure supplement 1). A Mann-Whitney non- parametric test was performed to detect statistically significant differences of XM and YM values.

### Statistical analysis

All statistical analysis and generation of graphs were performed using GraphPad Prism software (GraphPad Software Inc.). Data were subjected to a normality test to determine the statistical method to apply. Kruskal-Wallis followed by Dunn’s post-hoc test or Mann-Whitney were applied as non- parametric tests. Raw data and results of the statistical analysis are provided as source files linked to the corresponding figure.

### Transformation of *N. benthamiana* leaves, protoplast isolation and coimmunoprecipitation assay

Transgenic *Agrobacterium rhizogenes* (ARqua1) carrying the plasmid of interest were grown in liquid LB media supplemented with appropriate antibiotics for 24 hours at 28°C. Bacterial cultures were pelleted by centrifugation (4000 rpm), resuspended in Agromix (10 mM MgCl2; 10 mM MES/KOH pH 5.6; 150 uM Acetosyringone) and mixed in selected combinations to reach a final OD600 of 0.3. An *Agrobacterium tumefaciens* (GV3101) carrying a plasmid for expression of the silencing suppressor p19 was added to each mix (final OD600 = 0.1), which was then incubated at room temperature (RT) in the dark for 2 h. The second and third leaf of 4-5 weeks old *N. benthamiana* plants were infiltrated on the abaxial side using a 1 mL syringe. 72 h post-infiltration leaves were harvested and protoplasts were isolated as previously described (Su et al., 2022). Isolated protoplasts were directly lysed in 1 mL protein extraction buffer (50 mM Tris-HCl (pH 7.5), 150 mM NaCl, 10% (v/v) glycerol, 2 mM EDTA, 5 mM DTT, 1 mM phenylmethylsulfonyl fluoride (PMSF), Protease Inhibitor Cocktail (Roche), and 1% (v/v) IGEPAL CA-630 (I3021, Sigma-Aldrich)) and incubated for 1 h at 4°C. Lysates were centrifuged at 13000 rpm for 30 min (at 4°C) to obtain the supernatant fractions, which were then immunoprecipitated using RFP-Traps Magnetica agarose (rta, Chromotek) at 4°C for 90 min. After incubation with RFP-Traps, samples were washed 5 times with washing buffer (50 mM Tris-HCl (pH 7.5)), 150 mM NaCl, 10% (v/v) glycerol, 2 mM EDTA, Protease Inhibitor Cocktail (cOmplete^TM^, Mini, EDTA-free, 04693159001, Roche), and 0.5% (v/v) IGEPAL CA-630 (I3021, Sigma-Aldrich)). To release the proteins, 40 μL of washing buffer and 10 μL 5x protein loading buffer were added, and samples were heated for 10 min at 95°C. 10 μL of each sample were then loaded on a 12% (w/v) SDS gel, run for ∼90 mins (100 V, RT) and transferred overnight at 30 V (at 4°C) to a polyvinylidene difluoride (PVDF) membrane (Immobilon-P, pore size 0.45 μm, IPVH00010, Carl Roth). Transferred membranes were blocked with 5% (w/v) milk for 30 mins at RT before being hybridized with the corresponding anti-GFP (632381, Takara) or anti-mCherry (632543, Takara) antibodies at dilutions of 1:3000 and 1:2000, respectively, for 2 h at RT. Before incubating with the secondary antibody (anti-mouse, A-4415; anti-rabbit, A-6514; Sigma-Aldrich) (1 h, RT), membranes were washed 3 times for 10 min with TBST buffer. Membranes were developed following 3 washes of 10 min each with TBST buffer.

### Transmission electron microscopy

Wild-type (A17) and *rpg-1* germinated seedlings were transferred to an aeroponic system (25 liters volume), inoculated with *S. meliloti* (Sm2011, OD600=0.01, 5 l) and grown in liquid Fahräeus (0.1 mM NH4NO3) that was replaced weekly. Plants were harvested 35 dpi. Nodules were cut in half and immediately vacuum infiltrated for 15 min in MTSB buffer containing 4% PFA and 2.5% glutaraldehyde at room temperature. Samples were left in fixative for 3 h at room temperature and transferred to 4°C for overnight. Samples were washed with MTSB five times (10 min each) and post fixed in a water solution of 1% OsO4 on ice during 2.5 h. Samples were then washed with water twice for 10 min followed by *in bloc* staining with 1% UrAc in water for 1.5 h. After washing the samples twice with water for 5 min, they were subjected to dehydration by incubating them for 15 min each in increasing EtOH/water graded series (30%. 50%, 70%, 80%, 95%) and for 30 min twice in 100% EtOH and 100% Acetone. Samples were gradually embedded in Epoxi resin using resin:acetone mixtures (1:3, 1:1, 3:1) 8 h each, and finally pure resin (3 times exchange, 8 h each). Embedded nodules were polymerized for two days at 60°C and 70 nm sections were obtained with a Reichert- Jung Ultracut-E microtome. Sections were collected on copper grids and contrasted with 2% uranyl acetate and Reynolds lead citrate solution (Reynolds, 1963). Transmission electron microscopy images were obtained using a Philips CM10 (80 kV) microscope coupled to a GATAN Bioscan Camera Model 792.

## MATERIALS AVAILABILITY STATEMENT

All materials can be requested from the corresponding authors any time.

## ACKNOWLEDGEMENTS

We would like to thank Carla Brillada, Marco Trujillo and Franck A. Ditengou as well as Eija Schulze and Rosula Hinnenberg and Pascal Krohn for their experimental support and technical help, respectively, and the entire Ott lab team for the continuous inputs into the project. We would also like to thank Pierre-Marc Delaux and Jean Keller (LRSV, Université de Toulouse) for critically reading the manuscript and providing a comprehensive phylogeny on RPG, respectively, as well as Leonel Malacrida (Advanced Bioimaging Unit, IP Montevideo, Universidad de la República) for the fruitful discussions on Phasor analysis. We also thank the staff of the Life Imaging Center (LIC) in the Hilde Mangold House (HMH) of the Albert-Ludwigs-University of Freiburg for the help with their confocal microscopy resources, and the excellent support in image recording. The microscopes are operated by the Microscopy and Image Analysis Platform (MIAP) and the Life Imaging Center (LIC), Freiburg. TV belongs to the LRSV, which is part of the TULIP LABEX (ANR-10-LABX-41). The *Medicago truncatula* plants utilized in this research project, which are jointly owned by the Centre National De La Recherche Scientifique, were obtained from Noble Research Institute, LLC and were created through research funded, in part, by a grant from the National Science Foundation, NSF- 0703285.

## FUNDING

Engineering Nitrogen Symbiosis for Africa (ENSA) project currently supported through a grant to the University of Cambridge by the Bill & Melinda Gates Foundation (OPP1172165) and UK government’s Department for International Development (DFID) (TO)

Deutsche Forschungsgemeinschaft (DFG, German Research Foundation) 431626755 (TO)

DFG under Germany’s Excellence Strategy grant CIBSS – EXC-2189 – Project ID 39093984 (TO) China Scholarship Council (CSC) grants 201708080016 (CS)

DFG project number 414136422 (CLSM; TO), DFG project number 426849454 (TEM; TO)

## AUTHOR CONTRIBUTIONS

Conceptualization: BL, TO; Investigation: BL, CS, DIP, MR-F, TV, MB, SE, C-WL; Writing – Original Draft: BL, TO; Writing –Review & Editing: BL, CS, DIP, MR-F, TV, MB, SE, C-WL, TO; Supervision: BL, TO; Project administration: TO; Funding Acquisition: TO

---

**Figure 1 – figure supplement 1.**
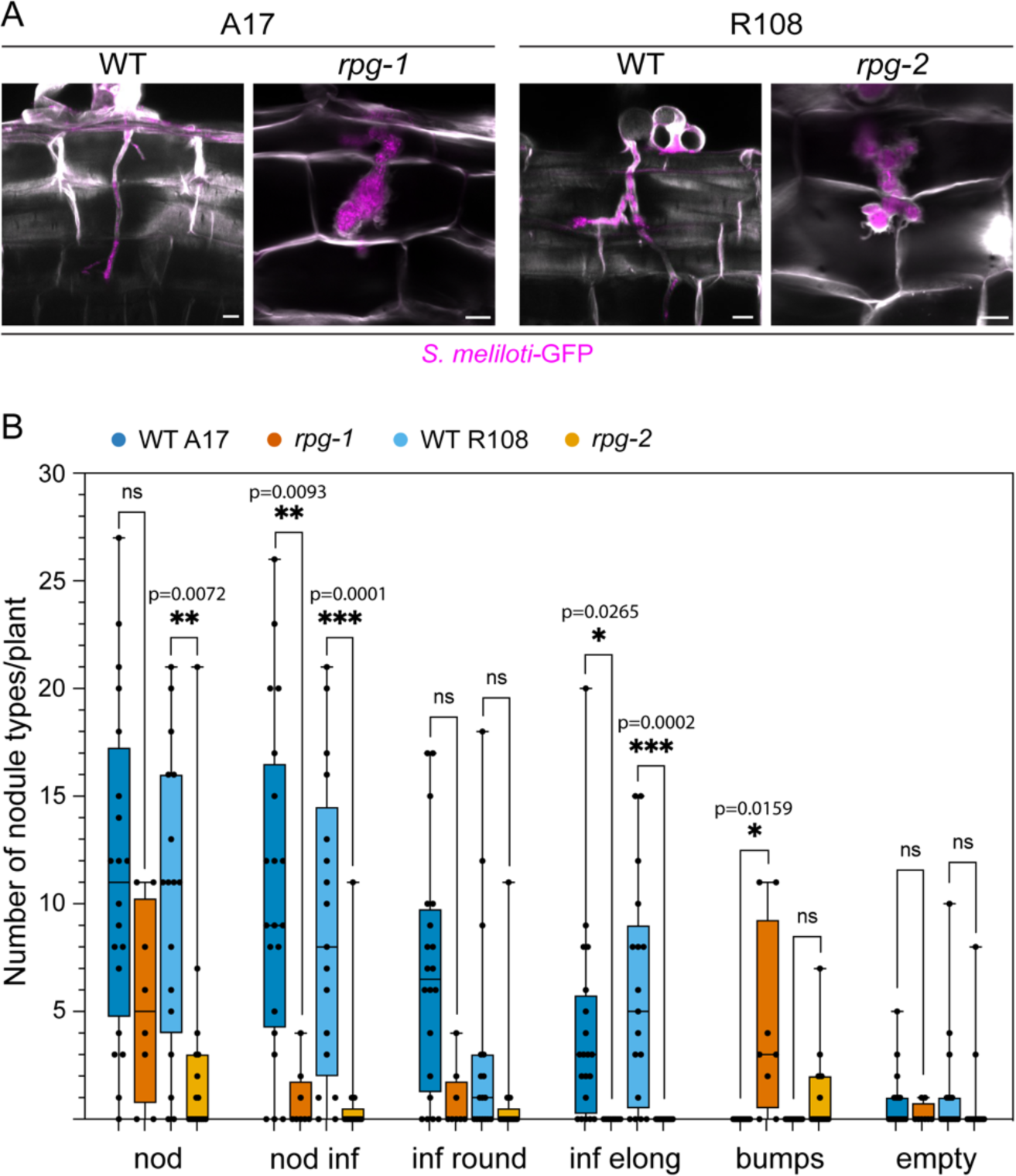
Cortical infection threads and nodule phenotype of *rpg* mutants. (**A**) Confocal images of cortical ITs formed in roots of *rpg* mutants compared to their corresponding wild-types (WT) at 14 dpi with *S. meliloti-GFP* (magenta). Cell walls were stained with Calcofluor white (white). Images are a merge of intensity projections of the fluorescent channels. Scale bars = 10 µm. (**B**) Number and type of nodules formed on *rpg* mutant alleles compared to their corresponding wild-types at 21 dpi with *S. meliloti*-LacZ. Data are presented as box plots where the top and bottom of each box represents the 75th and 25th percentiles, the middle horizontal bar indicates the median and whiskers represent the range of minimum and maximum values. Asterisks indicate statistical significance based on a Kruskal-Wallis multiple comparison analysis followed by Dunn’s post-hoc test with *p*-values <0.05 (*), <0.01 (**), < 0.001 (***). ns = not significant. The number of plants analyzed per genotype is 20 (WT A17), 8 (*rpg-1*), 17 (WT R108), 17 (*rpg-2*). Nod = total nodule structures; nod inf = total infected nodules; inf round = infected round nodules; inf elong = infected elongated nodules; bumps = bump-like nodules with arrested infection on the top; empty = empty nodules.

**Figure 2 – figure supplement 1.**
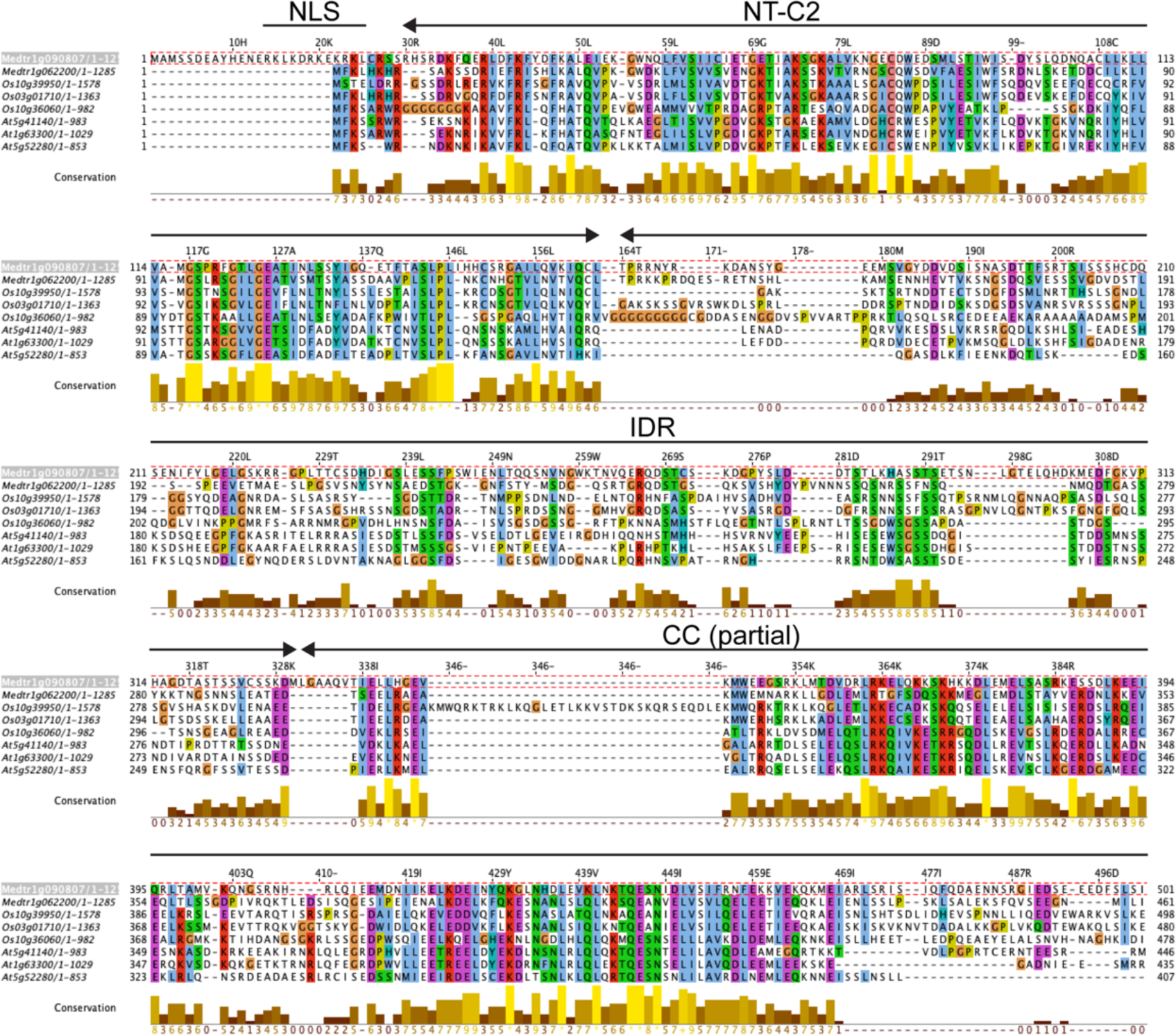
Sequence conservation among RPG homologs. Amino acid conservation between the NT-C2 domains and N-terminal parts of the coiled-coil region of RPG and homologous proteins from rice (Os) and Arabidopsis (At). Sequence alignment was performed using MUSCLE and visualized with Jalview to calculate conservation scores (Procter, 2021). Amino acids are colored according to Clustal X Colour Scheme. The sequence of *M. truncatula* RPG was used as reference for numbering of residues. NLS = nuclear localization signal; NT-C2 = N-terminal C2 domain; IDR = intrinsically disordered region; CC (partial) = coiled-coil region, partial.

**Figure 3-figure supplement 1.**
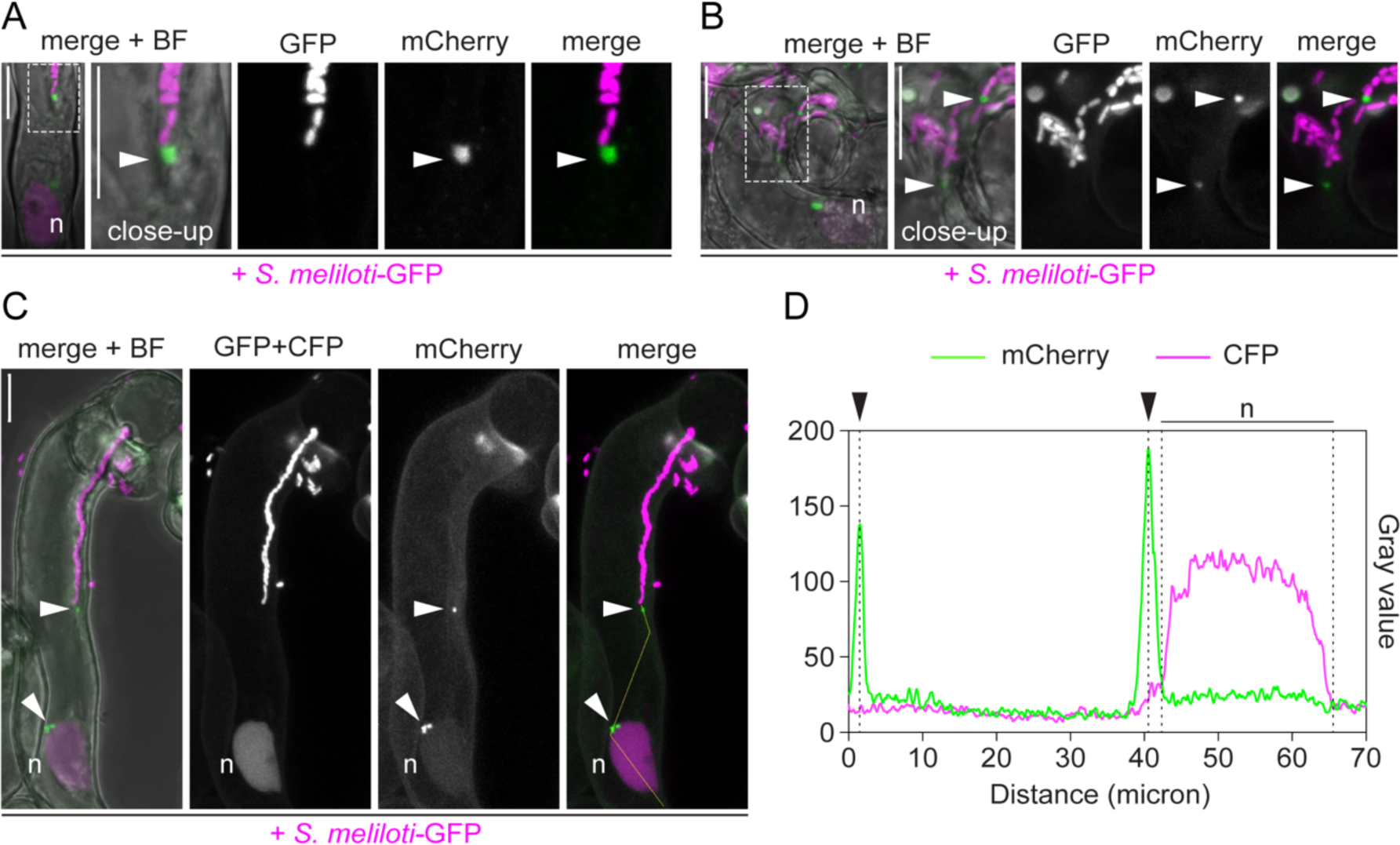
RPG sub-cellular localization pattern in root hairs. **(A-C)** Live-cell confocal imaging of infected root hairs of WT transgenic roots co-expressing mCherry-RPG and a nuclear localized tandem CFP as transformation marker. Images were acquired at 4-5 dpi with *S. meliloti*-GFP. **(A)** Punctate structure labelled by mCherry- RPG (arrowhead) capping the IT tip ahead of *S. meliloti*-GFP colonizing the tube. **(B)** mCherry-RPG puncta localize at the tip of ITs harboring multiple branches. **(C)** Aside with a bright signal located to puncta (arrowhead), a weak signal of mCherry-RPG can be occasionally observed in the nucleus (n) of infected root hairs. **(D)** Intensity profile along the yellow transect in **(C)**. Black arrowheads indicate intensity peaks on cytoplasmic puncta. The dashed white line boxes in **(A)** and **(B)** indicate the region shown in the respective close-ups. Individual channels are shown in grey; in merged images, the GFP/CFP channel is shown in magenta and the mCherry channel is shown in green. n = nucleus. Scale bars = 10 µm.

**Figure 3-figure supplement 2.**
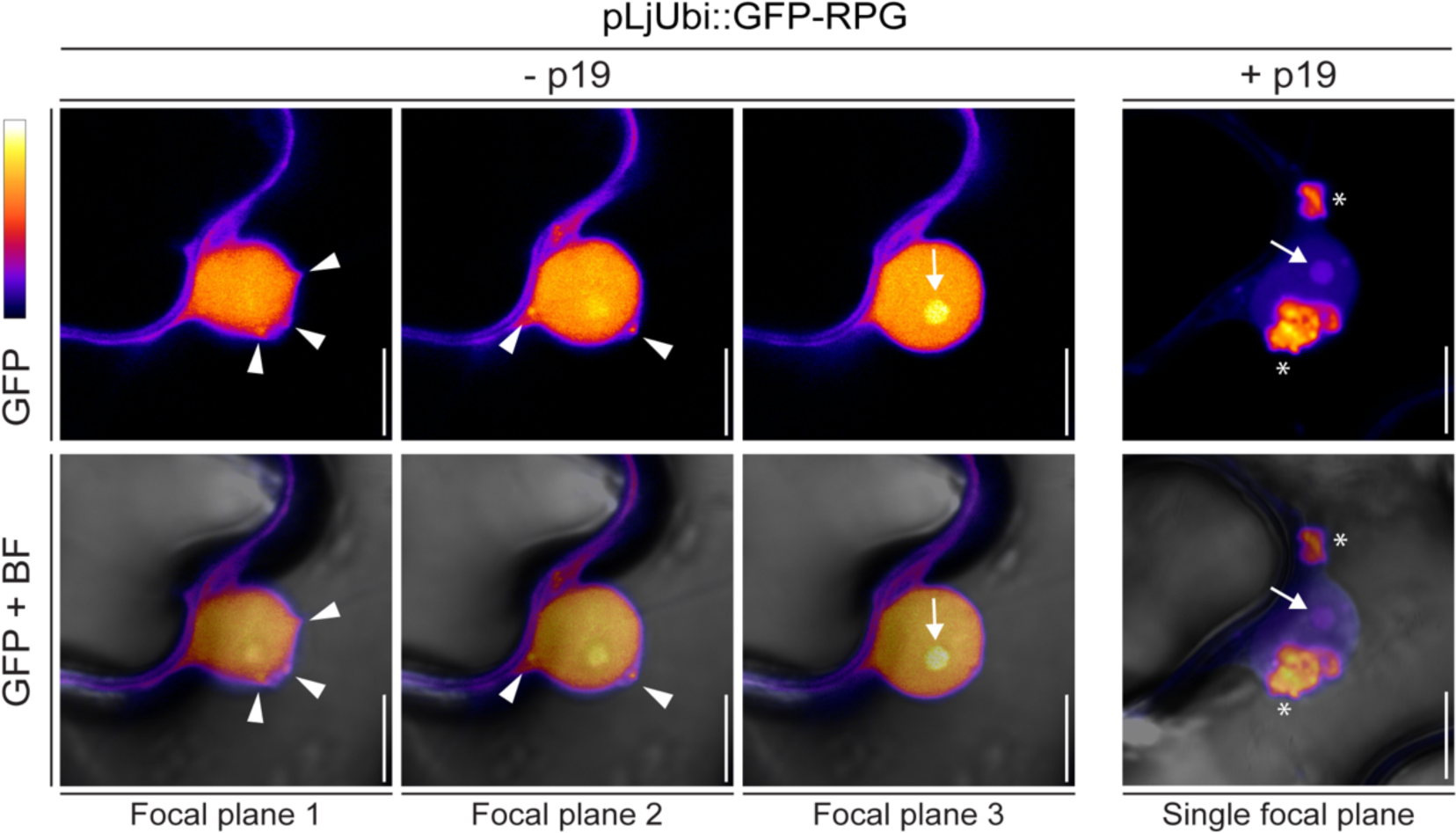
Subcellular localization of RPG in *N. benthamiana* epidermal cells. Live-cell confocal images of the nucleus and perinuclear region of *N. benthamiana* epidermal cells ectopically expressing GFP-RPG. In the absence of the silencing suppressor p19, GFP-RPG localizes in the nucleus, nucleolus (arrow) and in discrete puncta (arrowheads) located in the perinuclear region. Addition of p19 enhances accumulation of GFP-RPG in perinuclear associated structures (asterisks). The GFP channel is colored in Fire with yellow showing the maximum intensity and purple a low level of fluorescence. The bright field (BF) is overlaid with the GFP channel. Scale bars = 10 µm.

**Figure 3-figure supplement 3.**
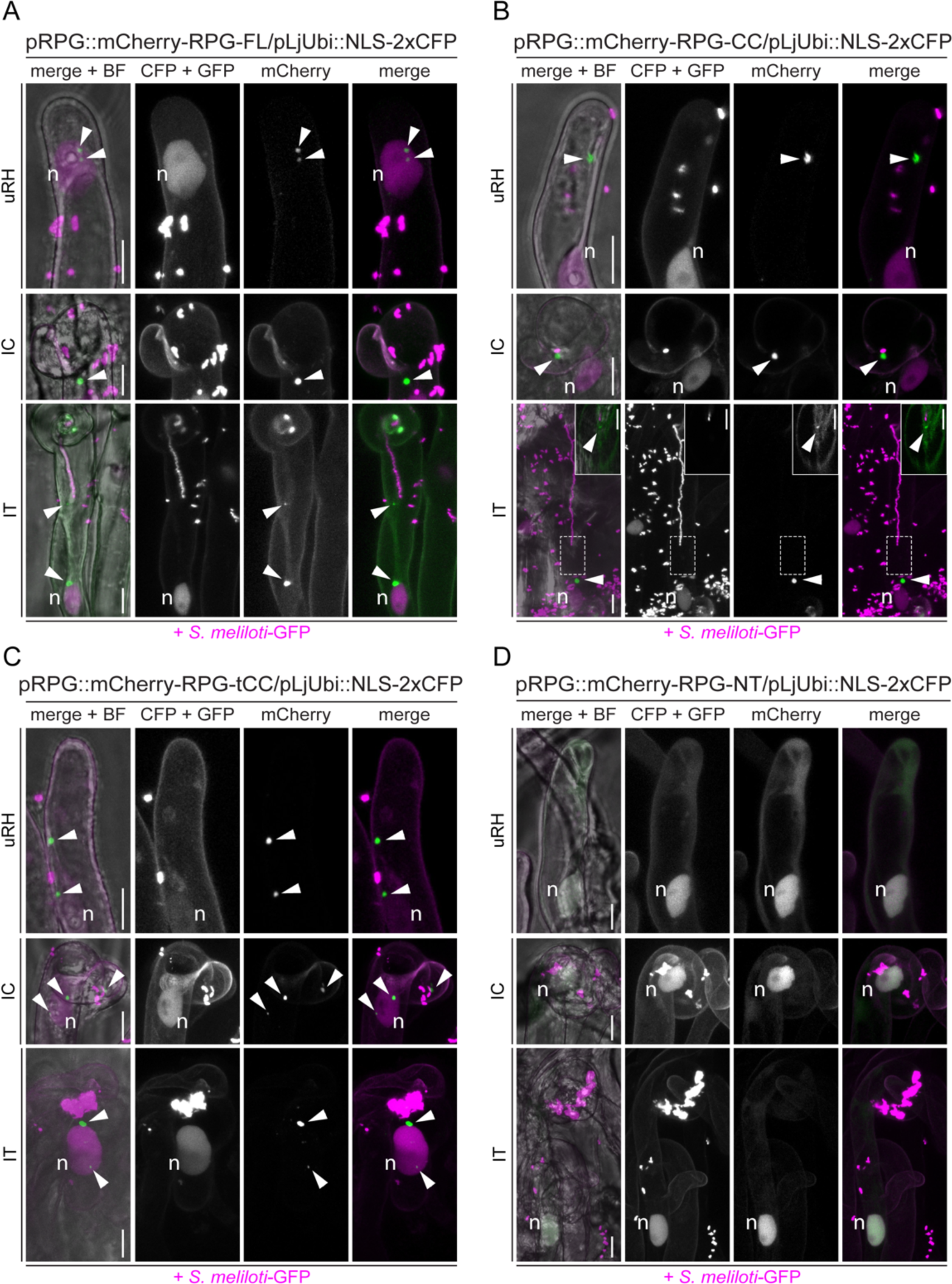
The coiled-coil region is necessary and sufficient to drive RPG localization to puncta. Live-cell confocal images of root hairs from *rpg-1* transgenic roots expressing mCherry-RPG and its deletion derivative imaged at 4-7 dpi with *S. meliloti*-GFP. For each construct, uninfected root hairs (uRH), curled root hairs hosting infection chambers (IC) and root hairs harboring infection threads (IT) were imaged. mCherry-FL **(A),** mCherry-RPG-CC **(B)** and mCherry-RPG-tCC **(C)** accumulate in bright punctate structures (arrowheads) in root hairs at different stages, while mCherry-NT **(D)** localizes in the nucleus and the cytoplasm. Images are intensity projections. Individual channels are shown in grey; in merged images, the GFP/CFP channel is shown in magenta and the mCherry channel is shown in green. A nuclear-localized tandem CFP was used as transformation marker. The dashed white line boxes in **(B)** indicate the region shown in the corresponding insets. Images in the insets are FLIM-based scans (20 repetitions) of single focal planes (CFP + GFP channel = 2.43 ns; mCherry channel = 1.38 ns; threshold = 10 photons). n = nucleus. Scale bars = 10 µm.

**Figure 3-figure supplement 4.**
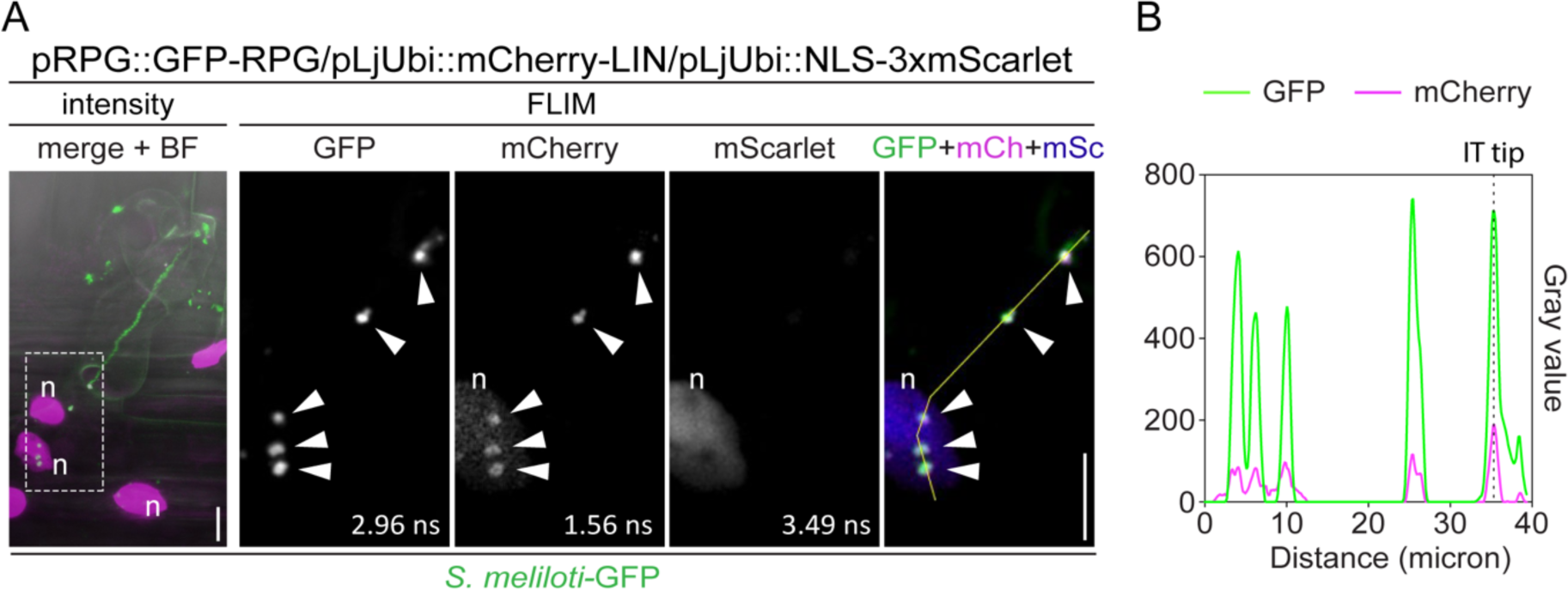
RPG co-localizes with LIN during root hair infection. **(A)** Live-cell confocal images of infected root hairs from transgenic WT roots showing co-localization of GFP-RPG with mCherry-LIN in punctate structures (arrowheads) located at the IT tip, in the perinuclear space and in the cytoplasm between these two poles. Weak signal of mCherry-LIN is visible in the nucleus (n). The intensity-based image is a merge of intensity projections of GFP (green) and mCherry/mScarlet (magenta) channels overlaid with the brightfield (BF) channel. The dashed white line box indicates the region shown in FLIM images. FLIM images are single focal planes with individual channels shown in grey. Lifetime values (ns) obtained from exponential reconvolution of decay profiles of GFP and mCherry/mScarlet channels are reported. When components are merged, GFP is shown in green, mCherry in magenta and mScarlet in blue. A nuclear-localized triple mScarlet was used as transformation marker. n = nucleus. Scale bars = 10 µm. Images are representative of 3 composite plants analyzed. **(B)** Intensity profile along the yellow transect shown in **(A)**. The intensity peaks at the infection thread tip (IT tip) are marked by a dashed black line.

**Figure 3-figure supplement 5.**
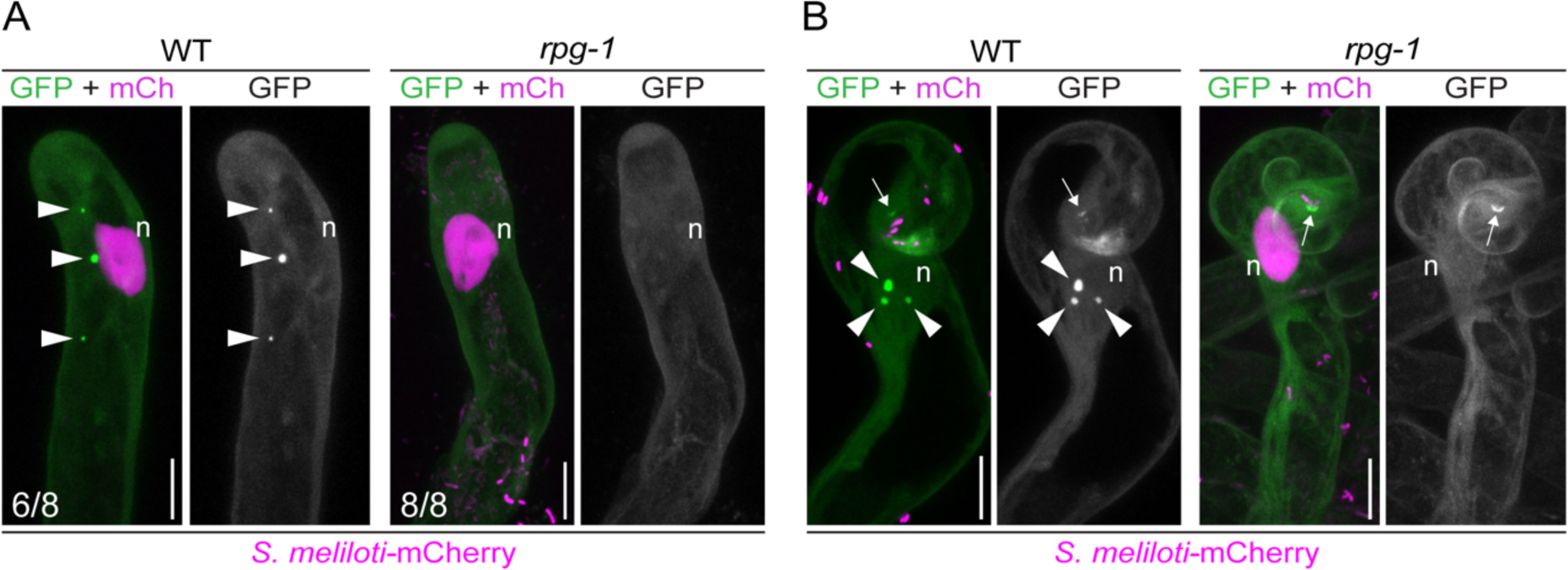
VPY recruitment to puncta is dependent on RPG. Live-cell confocal images of root hairs from transgenic WT and *rpg-1* roots expressing VPY-GFP. **(A)** VPY-positive cytoplasmic puncta (arrowheads) visible in WT uninfected root hairs are not maintained in *rpg-1.* **(B)** Accumulation of VPY-GFP adjacent to the site of rhizobia infection (arrows) occurs in few root hairs of *rpg-1* transgenic roots but it is not accompanied by bright punctate structures (arrowheads) associated to the nucleus (n) as seen in root hairs of WT transgenic roots. The GFP channel is shown in grey when isolated, in green when merged with mCherry (magenta). A nuclear-localized tandem mCherry was used as transformation marker. Numbers indicate frequencies of observations, made on a total number of 6 (WT) and 7 (*rpg-1*) composite plants. n = nucleus. Scale bars = 10 µm

**Figure 4-figure supplement 1.**
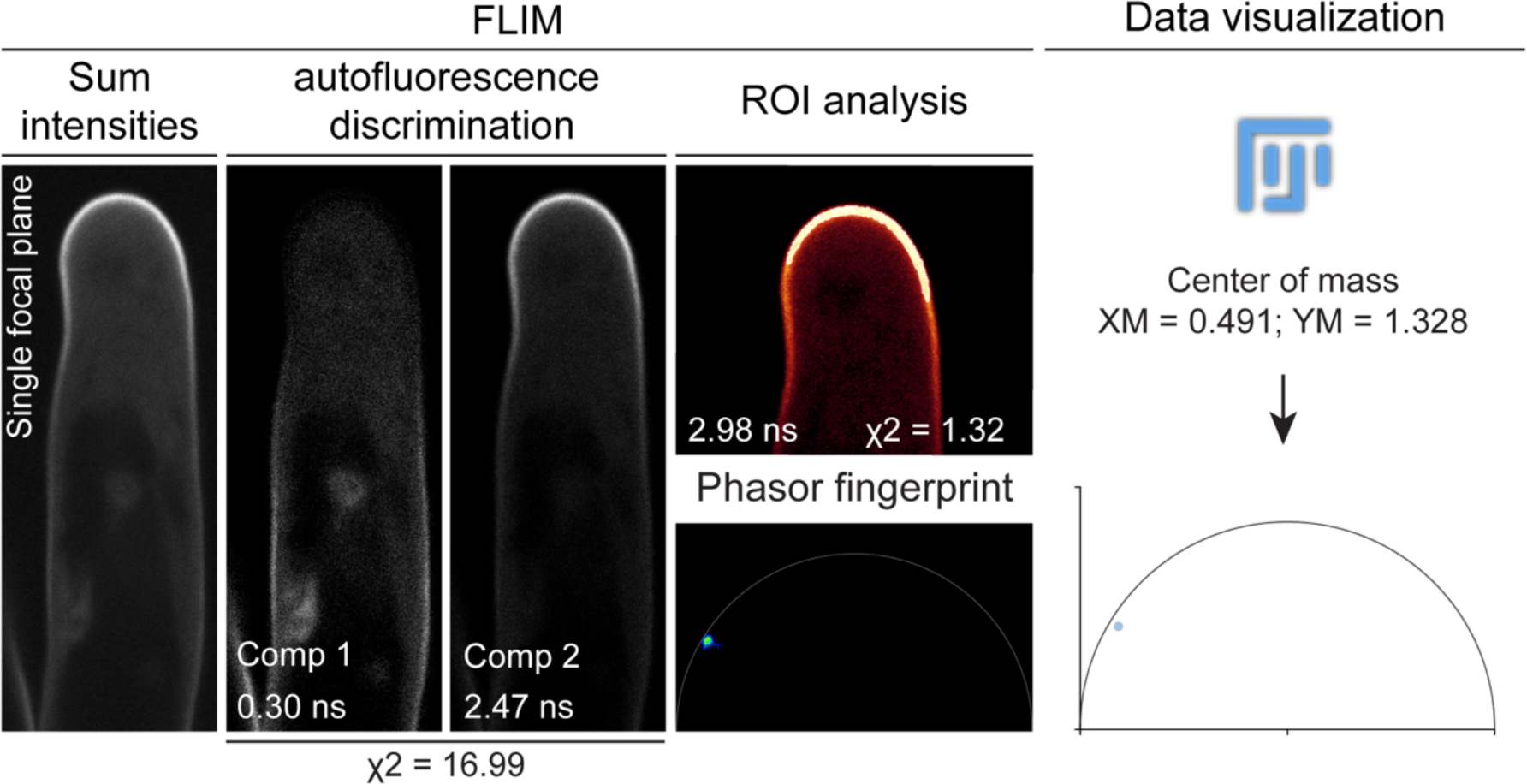
A FLIM-based systematic approach adopted to visualize PI(4,5)P2 membrane enrichment. The sequential steps used for visualization and analysis of PI(4,5)P2 distribution patterns are shown in a confocal image of a root hair from WT transgenic roots expressing mCitrine-PH^PLC^ as example. FLIM-based confocal imaging initially yields an image (sum intensities) where the intensities of the different fluorescent components present in the scanned area are summed up. By applying n-exponential reconvolution of the fluorescence intensity decay profile of the whole image, the major fluorescent components are separated and a lifetime value is assigned to each of them. The number of components is determined according to a reduced chi-squared (χ^2^) criterion (Lakowicz, 2006). The components exhibiting lifetime values ≤ 1.1 ns are considered as autofluorescence and are subtracted from the image. To determine with higher accuracy the enrichment of mCitrine-PH^PLC^ at the apical membrane domain, the decay profile at such region of interest (ROI) is analyzed. The lifetime is calculated via mathematical fitting, yielding a more precise estimation, as shown by the chi-squared (χ2) reaching values closer to 1. A lifetime value of ∼ 3 ns for mCitrine is consistent with previous reports (Söhnel et al., 2016). In addition, Phasor fingerprints of fluorescence emissions at the ROIs are generated using the fit-free Phasor approach, which assigns a position to each pixel in the ROI according to its composition in term of fluorescent species (Malacrida et al., 2021). Enrichment of the mCitrine-PH^PLC^ biosensor at the apical membrane domain results in pixels forming a cloud positioned on or close to the semicircle in the left part of the plot, where long mono-exponential decays are located. The Phasor fingerprint is further converted to a point in an XY plot by measuring the center of mass of the pixel population in the Phasor image using ImageJ/(Fiji) (Schindelin et al., 2012). In this way, data can be visualized on a graph.

**Figure 4-figure supplement 2.**
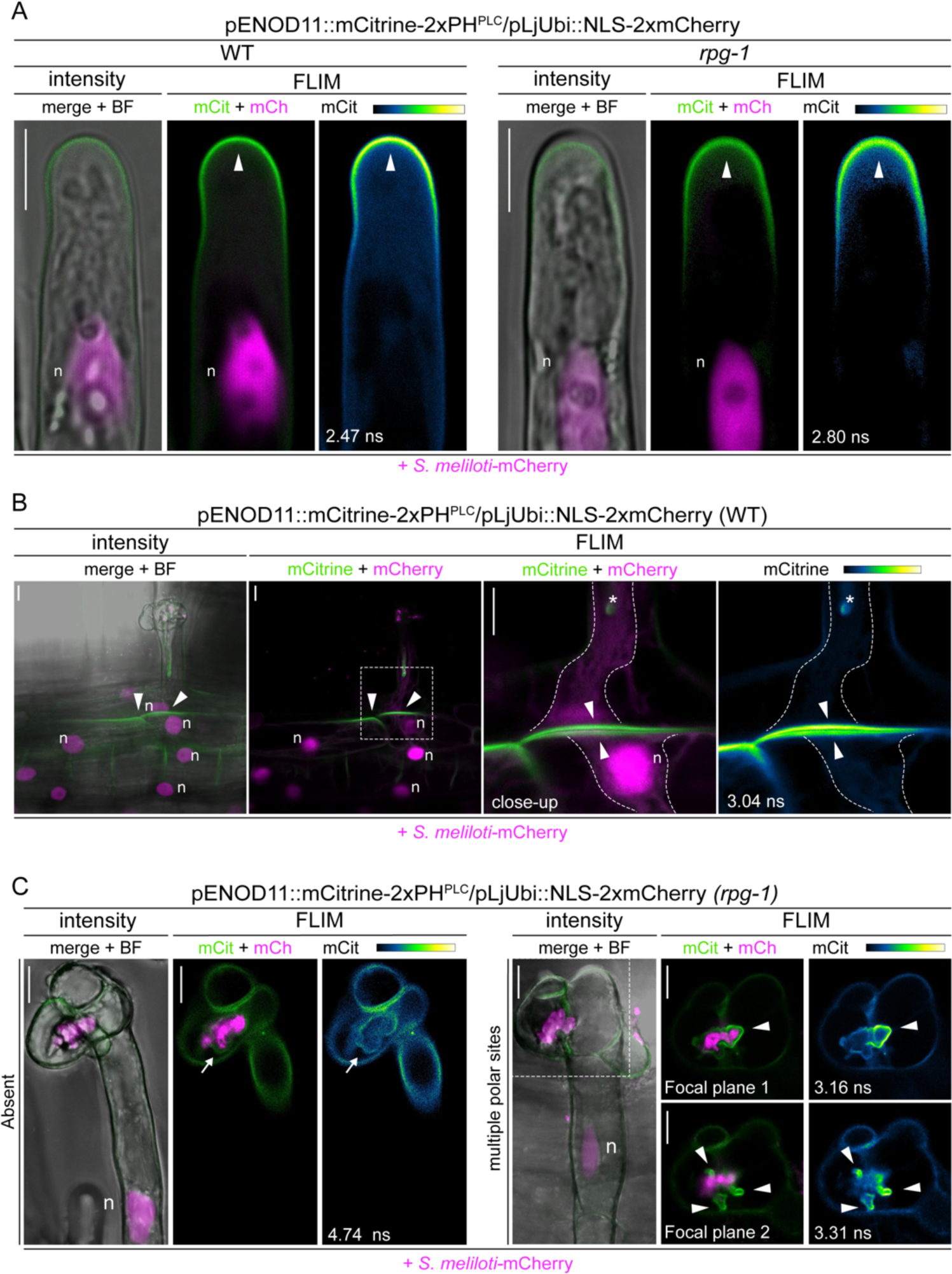
PI(4,5)P2 patterning in transgenic roots of WT and *rpg-1*. Live-cell confocal images of WT and *rpg-1* transgenic roots expressing the PI(4,5)P2 biosensor mCitrine-2xPH^PLC^ at 4-8 dpi with *S. meliloti*- mCherry. **(A)** Accumulation of PI(4,5)P2 (arrowhead) at the apical membrane of growing root hairs occurring in both WT and *rpg-1* transgenic roots. **(B)** PI(4,5)P2 enrichment (arrowheads) at the basal domain of infected root hairs and at the apical membrane of underlying cortical cells visualized in WT transgenic roots (enlarged panels of Figure 4A-CC). mCherry-derived cytoplasmic signals occurring due to leakage of the nuclear-localized transformation marker highlight the presence of a cytoplasmic column (dashed freehand line) connecting the tip of the IT (asterisk) to the base of the root hair and surrounding the nucleus (n) in the underlying cortical cells. The dashed white line box indicates the region shown in the close-up. **(C)** Representative images of infected root hairs on *rpg-1* transgenic roots showing an almost absent signal of the PI(4,5)P2 biosensor on the membrane (arrow, left panel) or its presence at multiple polar sites (arrowheads, right panel) on enlarged infection structures. The dashed white line indicates the region shown in focal plane 1 and 2. Intensity-based images are a merge of intensity projections of mCitrine (green) and mCherry (magenta) channels overlaid with the brightfield (BF) channel. FLIM images are single focal planes; the mCitrine component is shown in green when merged with the mCherry component (magenta) and in Green Fire Blue when isolated, with yellow indicating the maximum intensity and blue a low level of fluorescence. Lifetime values (ns) obtained from exponential reconvolution of decay profiles of the mCitrine channel are reported. n = nucleus. Scale bars = 10 µm.

**Figure 4-figure supplement 3.**
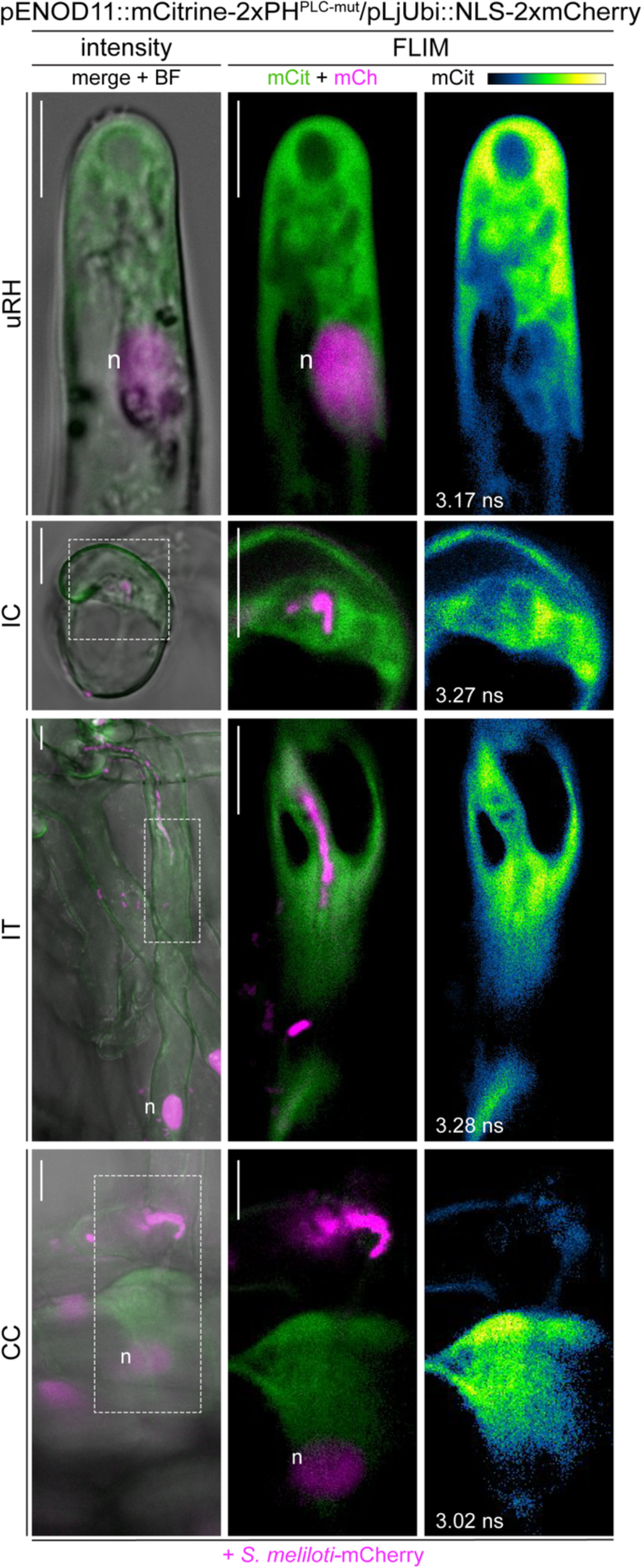
A mutant PI(4,5)P2 biosensor does not accumulate at membrane domains during rhizobial infection. Live-cell confocal images of WT transgenic roots expressing the mCitrine-2xPH^PLC^ ^mut^ biosensor at 4-7 dpi with *S. meliloti*-mCherry. The signal from the mutated version of the biosensor appeared diffused in the cytoplasm of uninfected (uRH) and infected (IC-IT) root hairs, and in cortical cells (CC) on the infection thread trajectory. Intensity-based images are a merge of mCitrine (green) and mCherry (magenta) channels overlaid with the brightfield (BF) channel. FLIM images are single focal planes; the mCitrine component is shown in green when merged with the mCherry component (magenta) and in Green Fire Blue when isolated, with yellow indicating the maximum intensity and blue a low level of fluorescence. Lifetime values (ns) obtained from exponential reconvolution of decay profiles of the mCitrine channel are reported. Images are representative of 3 composite plants analyzed. n = nucleus. Scale bars = 10 µm.

**Figure 4-figure supplement 4.**
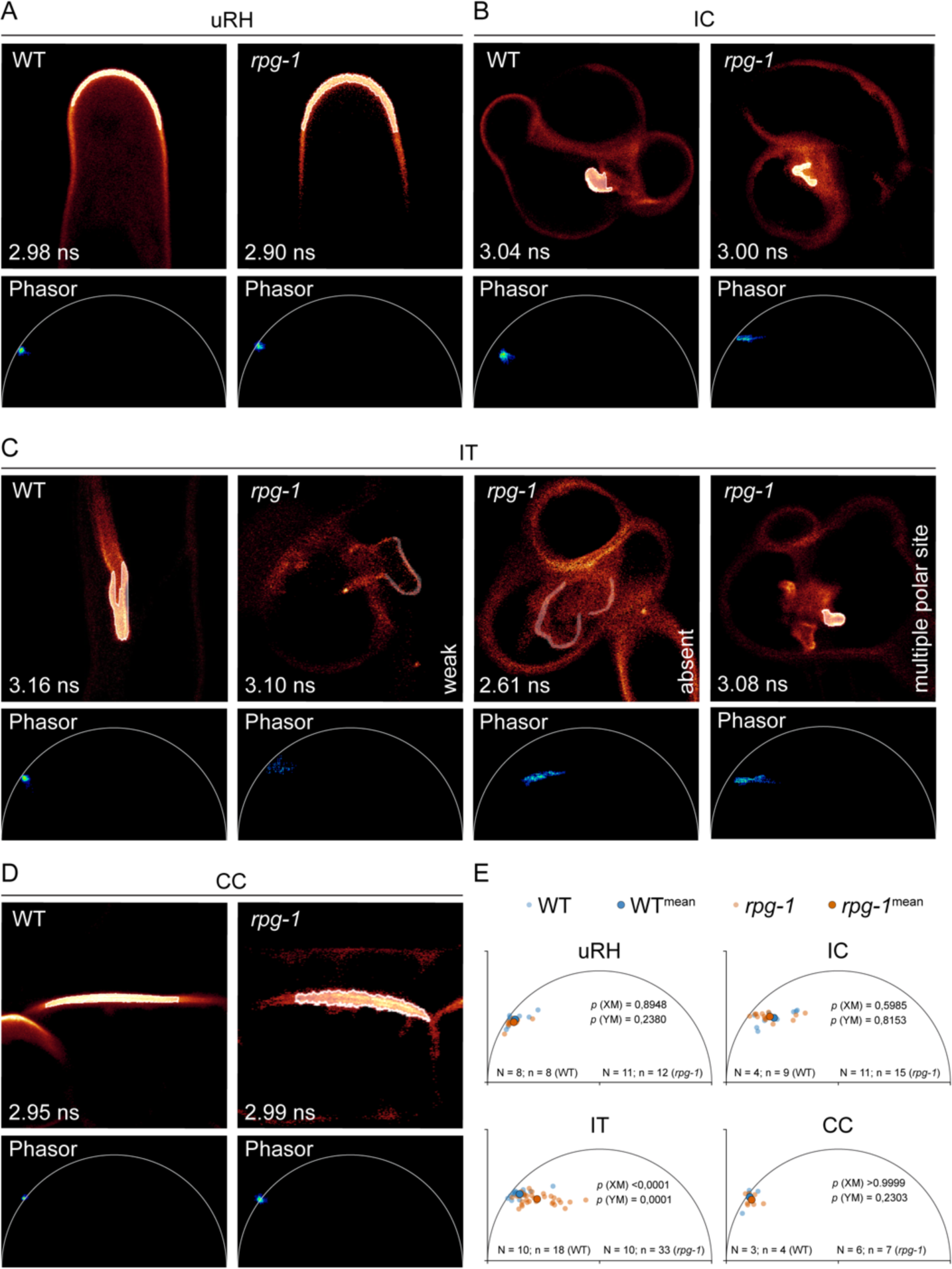
Phasor analysis statistically supports loss of IT membrane polarization occurring in *rpg-1.* Estimation of PI(4,5)P2 membrane enrichment in WT and *rpg-1* transgenic roots expressing the PI(4,5)P2 biosensor mCitrine-PH^PLC^ via analysis of fluorescence emission at regions of interest (ROI) corresponding to different membrane domains. **(A-D)** Representative FLIM images of analyzed membrane domains showing the Phasor fingerprint (Phasor) associated to the ROI selected on the membrane (white outlines) and the lifetime value (ns) calculated via fitting of the decay profile of the ROI. Phasor fingerprints of ROIs selected at the apex of uninfected root hairs (uRH) **(A)**, at the site of symbiont emergence from the infection chamber (IC) **(B)** and on the membrane domain on the trajectory of IT propagation in cortical cells (CC) **(D)** in cells of WT and *rpg-1* transgenic root show similar positioning on the left half of the Phasor plot close to the semicircle, indicative of similar PI(4,5)P2 enrichment. **(C)** The Phasor signature associated to the membrane of ITs in *rpg-1* markedly differs from the signature of WT ITs: a pixel cloud with similar Phasor position but composed of few scattered pixels is indicative of weak signal of the mCitrine-PH^PLC^ biosensor, while a consistent shift of the pixel cloud away from the semicircle towards the right part of the plot where short lifetimes are located is indicative of dominance of autofluorescent components over mCitrine-PH^PLC^, whose signal appears mostly absent. The Phasor fingerprint associated to the membrane surrounding the multiple polar sites emerging from the enlarged infection threads in *rpg-1* is similar to the one recorded at the IC of *rpg-1* in **(B)**. **(E)** XY plots representing the center of mass calculated from Phasor fingerprints of different membrane domains analyzed in multiple cells of different plants for each genotype. A Mann-Whitney test was applied to detect statistical differences between XM and YM values. Calculated *p* values are shown. N = number of plants; n = number of cells.

**Figure 4-figure supplement 5.**
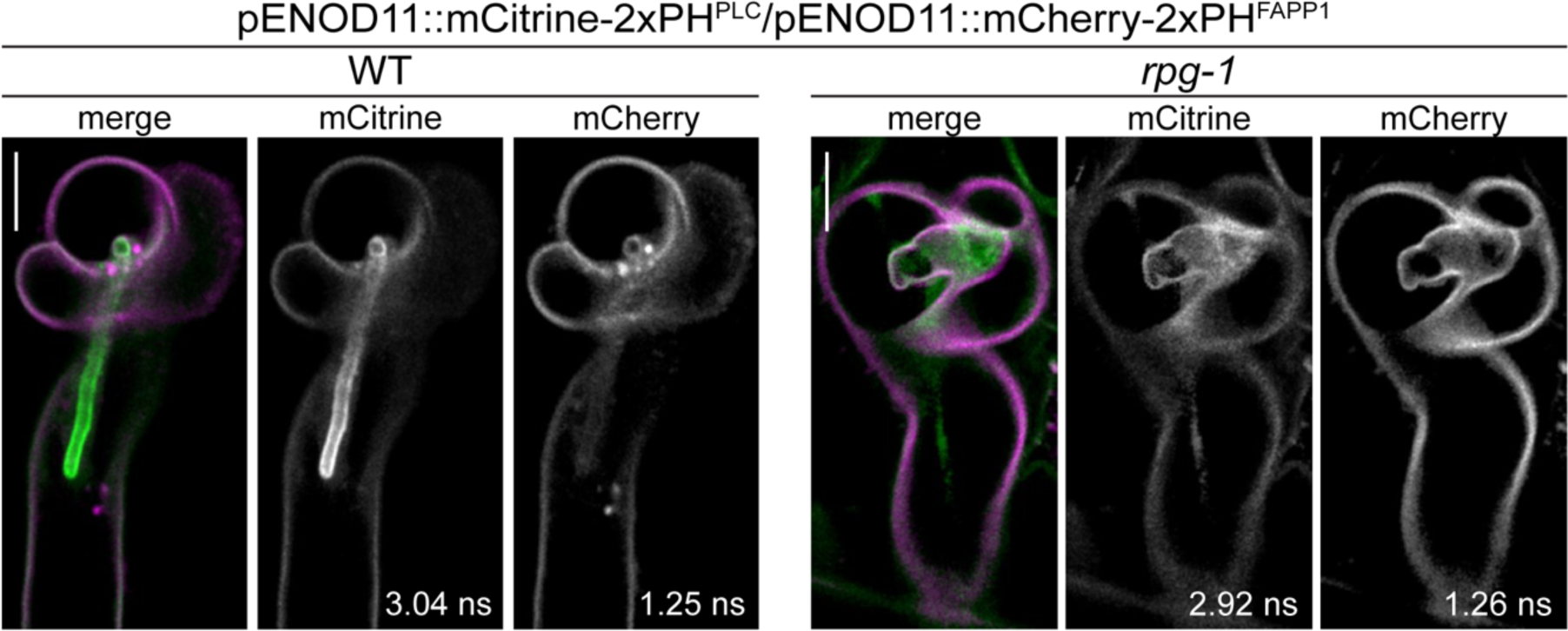
Co-visualization of PI4P and PI(4,5)P2 in WT and *rpg-1* infected root hairs. FLIM-based confocal images of infected root hairs shown in Figure 4B from WT and *rpg-1* transgenic plants co-expressing mCitrine-2xPH^PLC^ and mCherry-2xPH^FAPP1^ at 4 dpi with *S. meliloti*-LacZ. Images are single focal planes with individual channels shown in grey. Lifetime values (ns) obtained from exponential reconvolution of decay profiles of mCitrine and mCherry channels are reported. When components are merged, mCitrine is shown in green and mCherry in magenta. Scale bars = 10 µm.

## Source Data Files

Figure 1 – source data 1

IT diameter and statistical analysis

Figure 1 – figure supplement 1 – source data 1

Nodule number and statistical analysis

Figure 2 – source data

IT diameter and statistical analysis

Figure 3 - source data 1

FLIM fit values and intensity profile values

Figure 3 - source data 2

Raw images of Western Blot analysis

Figure 3 – figure supplement 1 – source data 1

Intensity profile values

Figure 3 – figure supplement 4 – source data 1

FLIM fit values and intensity profile values

Figure 4 - source data 1

FLIM fit values and intensity profile values

Figure 4 – figure supplement 1 – source data 1

FLIM fit values

Figure 4 – figure supplement 2 – source data 1

FLIM fit values

Figure 4 – figure supplement 3 – source data 1

FLIM fit values

Figure 4 – figure supplement 4 – source data 1

FLIM fit values and Phasor analysis values

